# Fe starvation induces a second LHCI tetramer to photosystem I in green algae

**DOI:** 10.1101/2024.12.11.624522

**Authors:** Helen W. Liu, Radhika Khera, Patricia Grob, Sean D. Gallaher, Samuel O. Purvine, Carrie D. Nicora, Mary S. Lipton, Krishna K. Niyogi, Eva Nogales, Masakazu Iwai, Sabeeha S. Merchant

## Abstract

Iron (Fe) availability limits photosynthesis at a global scale where Fe-rich photosystem (PS) I abundance is drastically reduced in Fe-poor environments. We used single-particle cryo-electron microscopy to reveal a unique Fe starvation-dependent arrangement of light-harvesting chlorophyll (LHC) proteins where Fe starvation-induced TIDI1 is found in an additional tetramer of LHC proteins associated with PSI in *Dunaliella tertiolecta* and *Dunaliella salina*. These cosmopolitan green algae are resilient to poor Fe nutrition. TIDI1 is a distinct LHC protein that co-occurs in diverse algae with flavodoxin (an Fe-independent replacement for the Fe-containing ferredoxin). The antenna expansion in eukaryotic algae we describe here is reminiscent of the iron-starvation induced (*isiA-*encoding) antenna ring in cyanobacteria, which typically co-occurs with *isiB*, encoding flavodoxin. Our work showcases the convergent strategies that evolved after the Great Oxidation Event to maintain PSI capacity.

## Main text

About half of Earth’s primary productivity occurs through photosynthesis by marine algae (*1*). The photosynthetic machinery evolved in an Fe-rich anoxic world, but in today’s oxygenated world, Fe-bioavailability is poor due to the low solubility of ferric species (*2*, *3*). Three multi-subunit membrane protein complexes in photosynthesis, photosystem (PS) II, the cytochrome (Cyt) *b*_6_*f* complex, and PSI, catalyze sequential electron transfer from water to generate reductant for biosynthesis, and all require Fe as a cofactor. These complexes are a major Fe sink in the cell given their intracellular abundance and their high Fe stoichiometry (*4–6*). PSI has the highest stoichiometry, with three iron-sulfur (Fe_4_S_4_) clusters per monomer, corresponding to about half the Fe in the photosynthetic apparatus, and it is thus disproportionately impacted by poor Fe nutrition. Consequently, the PSI content of green algae, diatoms and cyanobacteria is reduced substantially under Fe starvation, with an impact on photosynthetic productivity and biomass. But little else is known about the molecular mechanisms of algal PSI adaptations to low Fe conditions in the ocean (*7–9*).

To address this gap, we surveyed the globally abundant, cosmopolitan green algae, *Dunaliella* spp. (Fig. 1a). These algae are obligate photoautotrophs, that tolerate extreme pH, salinity, light, and temperature, making them compelling organisms to investigate the diversity of both photoacclimation and Fe starvation response mechanisms (*10*). *Dunaliella* spp. are well- adapted to low Fe conditions, maintaining photosynthesis at Fe concentrations that are limiting to other photosynthetic organisms (Extended Data Fig. 1) (*10–13*). Here, we used two halotolerant *Dunaliella* spp. that diverged ∼253 million years ago: *D. tertiolecta* (isolated from the coastal Arctic seawater environment of Oslofjord, Norway) and *D. salina* (isolated from hypersaline Lake Bardawil in Egypt) (Fig. 1a) (*13*). Both organisms have an expanded repertoire of Fe homeostasis genes and express a range of similar Fe starvation responses that enables productivity under Fe- poor conditions, including upregulation of Fe-acquisition pathways, replacement of Fe-containing ferredoxin with flavin-containing flavodoxin, and downregulation of the Fe-containing photosynthetic apparatus, especially PSI components (*11*, *13–17*).

**Fig. 1.**
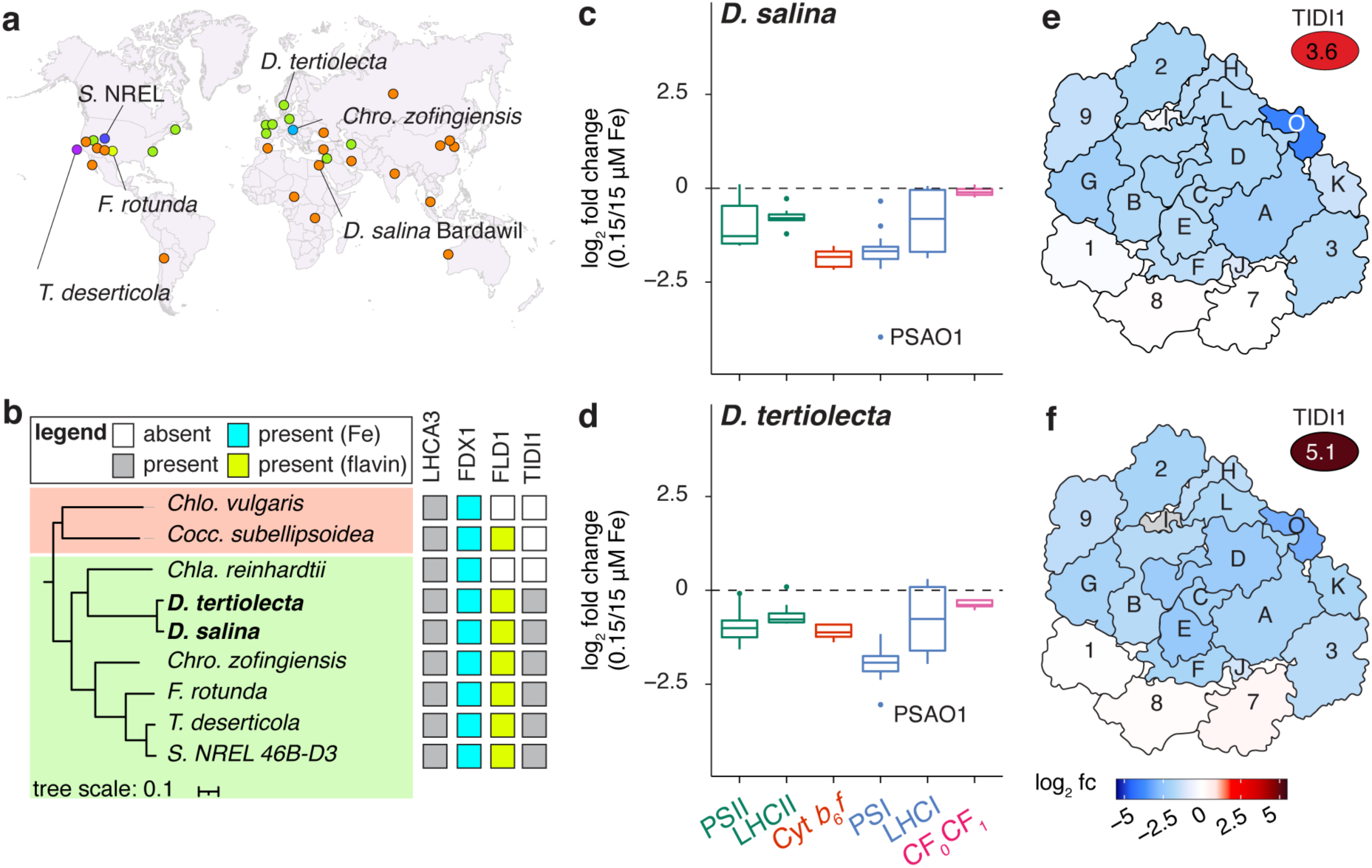
Uneven response of *Dunaliella* spp. LHCA subunits in Fe-starved medium. **a**, Geographic distribution of TIDI-encoding algae of either *D. salina* (orange), *D. tertiolecta* (green), *Chro. zofingiensis* (dark blue), *T. deserticola* (purple), *F. rotunda* (yellow), and *S. NREL 46B-D3* (light blue) modified from Davidi et al. (2024). The strains used in this work are *D. tertiolecta* UTEX LB999 and *D. salina* Bardawil UTEX LB 2538. **b**, A species tree showing the relationship between TIDI1 and non-TIDI1 encoding species from Chlorophyceae (green) and Trebouxiophyceae (salmon). The tree is based on the concatenated polypeptide sequences of universal single-copy orthologs (USCOs) shared among all 9 species. Filled squares signifies the presence of at least one homolog of the indicated protein (FDX1, ferredoxin; FLD1, flavodoxin; LHCA3; TIDI1). Square color describes proteins that are either present (gray), present and binds Fe (cyan), present and binds flavin (green), or absent (white). **c**,**d**, The log_2_ fold changes (fc) of protein abundances in the Fe-replete (15 µM Fe) and Fe-starved (0.15 µM Fe) condition detected by at least two spectral counts encoding components of PSII, LHCII, Cyt *b*_6_*f*, PSI, LHCI, and CF_O_CF_1_ (ATP synthase) for *D. salina* (**c**) and *D. tertiolecta* (**d**). For a complete list of protein subunits and their abundances, see Extended Data 2. **e**,**f**, The PSI (PSA) and LHCI (LHCA) subunits of PDB:6SL5 (stromal view) colored by its log_2_ fold changes (fc) of protein abundances (red, increase; blue, decrease) for *D. salina* (**e**) and *D. tertiolecta* (**f**). Gray values indicate not detected (<2 spectral counts).

In contrast to the decreased expression of PSI subunits, the expression of one thylakoid membrane protein, named TIDI1 (thylakoid iron deficiency induced), is dramatically increased under Fe starvation, similarly to the expression of FLD1, a flavodoxin (Extended Data Fig. 1, g and h) (*13*, *18*). A survey of green algal genomes found TIDI1 orthologs in a subset of green algae: *Chromochloris zofingiensis*, *Flechtneria rotunda*, *Tetradesmus deserticola*, and *Scenedesmus* sp. NREL 46B-D3 (*13*) (Fig. 1b). These algae were isolated from diverse habitats, including fresh water, brackish waters, and the desert, indicating that TIDI1 is neither a species- specific nor an environment-specific adaptation (*10*, *19*, *20*) (Fig. 1, a and b). As an intriguing correlation, each of these organisms also contains an ortholog of eukaryotic FLD1, consistent with the co-function of TIDI1 and FLD1 in a low Fe environment (*13*, *21*). Sequence analysis places TIDI1 in the family of light-harvesting chlorophyll (LHC) proteins (*12*, *13*, *22*). TIDI1 shares ∼50% sequence similarity to LHCA3s, which occur universally in the green lineage (*12*, *13*, *22*). LHCA3 binds to PSI on the PsaA/PsaK1 side and is on one path of energy transfer from the accessory antenna to the core antenna (*23*). Among its unique features, TIDI1 has a proline-rich stromal N-terminal domain and a longer lumenal loop between the transmembrane helices B and C (*13*). TIDI1 co-migrates with PSI-LHCI supercomplexes from Fe-starved *D. tertiolecta*, suggesting that TIDI1 may function in an Fe starvation-modified antenna system (*12*). However, the nature of the association and the placement of TIDI1 within the green algal antenna are not known.

In this work, we report high-resolution cryo-electron microscopy (cryo-EM) structures of PSI-LHCI supercomplexes from Fe-starved compared to Fe-replete cells of both *D. salina* and *D. tertiolecta*. We discovered that the PSI-LHCI supercomplexes under Fe-starved conditions have an additional LHCI tetramer in which TIDI1 replaces LHCA3 in the new tetramer. These structures represent new examples of LHC arrangements and illustrate the diversity of PSI-LHCI supercomplexes among photosynthetic organisms. This work constitutes the first structural depiction of eukaryotic PSI-LHCI antenna remodeling as an adaptive mechanism in response to Fe starvation.

### LHCA1, LHCA7, and LHCA8 are selectively maintained in Fe-starved medium

To quantify the changes in abundance of the proteins of the photosynthetic machinery under Fe starvation, we performed tandem mass tag (TMT) quantitative proteomics using cultures grown in Fe-replete (15 µM Fe) and Fe-starved (0.15 µM Fe) conditions. Principal component analysis (PCA) of the entire proteome revealed that >75% of the variance in protein abundance is captured in the first two components for both *D. salina* and *D. tertiolecta*, and most of the variance (>64%) is attributed to Fe availability (Extended Data Fig. 2, a and b). Apart from ATP synthase (CF_0_CF_1_), whose function is independent of Fe and whose abundance is unchanged, each of the components of the photosynthetic machinery decreased in abundance, albeit not to the same extent (Fig. 1, c and d). The most dramatic changes are for PSI, whose subunits decrease about 4-fold relative to the Fe-replete *Dunaliella* cells. This result is not unexpected given PSI’s high Fe content. By contrast, PSII, which requires only two Fe atoms per core monomer, showed less decrease (Fig. 1, c and d). While most of the PSI subunits (denoted by PSA in their names) decreased equivalently in low Fe medium, PSAO1 (known to be important for state transitions in *Arabidopsis thaliana* (*24*)) showed a more substantial decrease of 16- and 8-fold in *D. salina* and *D. tertiolecta*, respectively.

PSI and PSII are each associated with LHC proteins for increased absorption cross-section, LHCA for PSI and LHCB (or LHCBM) for PSII. These proteins are organized in the LHCI and LHCII oligomeric complexes, respectively. When we monitored the abundances of LHCI and LHCII subunits, we noted that the six *Dunaliella* LHCA proteins (numbered according to *Chlamydomonas reinhardtii* LHCA orthologs) are, like the PSA subunits, more substantially reduced in low Fe relative to the PSII subunits (PSB) and LHCB proteins (Fig. 1, c and d). Nevertheless, quantitative proteomics data indicated a striking difference between the abundance of three LHCA proteins (LHCA1, LHCA8 and LHCA7), which remain unchanged, and the other LHCA proteins (LHCA2, LHCA3 and LHCA9), which decreased proportionately with PSI subunits, as expected (Fig. 1, e and f). TIDI1, on the other hand, was greatly increased in low Fe medium in both *Dunaliella* spp., as previously shown (*12*, *13*). The changes in protein abundance mirror changes in transcript abundances (*13*), indicative of an antenna remodeling program in Fe starvation (Fig. 1, e and f). These results motivated structural analysis of PSI-LHCI supercomplexes.

### Isolation of Dunaliella PSI-LHCI supercomplexes from Fe-replete and Fe-starved cells

We isolated PSI-LHCI supercomplexes from Fe-replete (15 µM Fe) and Fe-starved (0.15 µM Fe) cultures of both *Dunaliella* spp. (see Methods and Extended Data Fig. 3, a and d) using density gradient centrifugation of detergent-solubilized thylakoid membranes. The PSI supercomplexes from Fe-starved cells appeared in denser fractions of the gradient compared to those from Fe-replete cells (Extended Data Fig. 3, a and d), with *D. salina* separating in even denser fractions relative to *D. tertiolecta*, intimating larger PSI-LHCI supercomplexes in the Fe-starved situations. Immunoblot analysis indicated the presence of TIDI1 in these denser fractions (Extended Data Fig. 3, b and e). Low-temperature fluorescence emission spectra of each PSI-containing fraction exhibited a single peak around 710 nm, confirming the purity of the isolated PSI-LHCI supercomplexes, with the peak from the Fe-starved cultures being 2-3 nm blue-shifted (Extended Data Fig. 3, c and f), perhaps reflecting a change in PSI antenna organization (*25*).

### Overall structures of Dunaliella spp. PSI-LHCI supercomplexes from Fe-replete cells

We used single-particle cryo-EM to determine the structures of both *D. salina* (*Ds*) and *D. tertiolecta* (*Dt*) PSI-LHCI supercomplexes from Fe-replete cultures as exemplars of the native state (hereafter named PSI-LHCI_1_) at overall resolutions of 2.9 Å and 2.1 Å, respectively (See Methods and Fig. 2, a and c, Extended Data Fig. 4, 5, and 6, Extended Table 1). The density maps were better defined around the PSI core than near the detergent belt (Extended Data Fig. 4, 5, and 6), consistent with a higher level of flexibility in the outer regions of the complex. Three sub-complex features are evident in the structures: the PSI core, consisting of all but one of the 12 PSA subunits encoded in the genome (PsaA to PSAL1); one crescent-shaped LHCI tetramer made up of four LHCA subunits (LHCA1, LHCA8, LHCA7 and LHCA3) arranged from the PSAG1 (proximal) to the PSAK1 (distal) side; and one LHCI dimer (LHCA2 and LHCA9) on the opposite side, together accounting for all 6 LHCA proteins encoded in the genome (Extended Table 2). We lacked density for the PSAO1 subunit, which may be attributed to our isolation or vitrification conditions (Fig. 2a, Extended Data Fig. 6, (*26*)), since the protein is detected in whole cell extracts (Fig. 1, e and f) albeit with undetermined occupancy in the PSI-LHCI supercomplex. Among the ∼300,000 particles evaluated, we did not find a previously reported *D. salina* “minimal” PSI-LHCI supercomplex lacking the LHCI dimer, PSAG1, PSAH1, PSAI1, PSAL1, and PSAO1 (Extended Data Fig. 4 and 5) (*27*). Otherwise, our structure recapitulates the more typical PSI supercomplex structure (*26*). The coordinate decrease of all PSA subunits in Fe-starved cells, evident in whole-cell proteomics data (Fig. 1, e and f, and Extended Dataset 1 and 2), argues in favor of the more complete PSI structure as the predominant *in vivo* species.

**Fig. 2.**
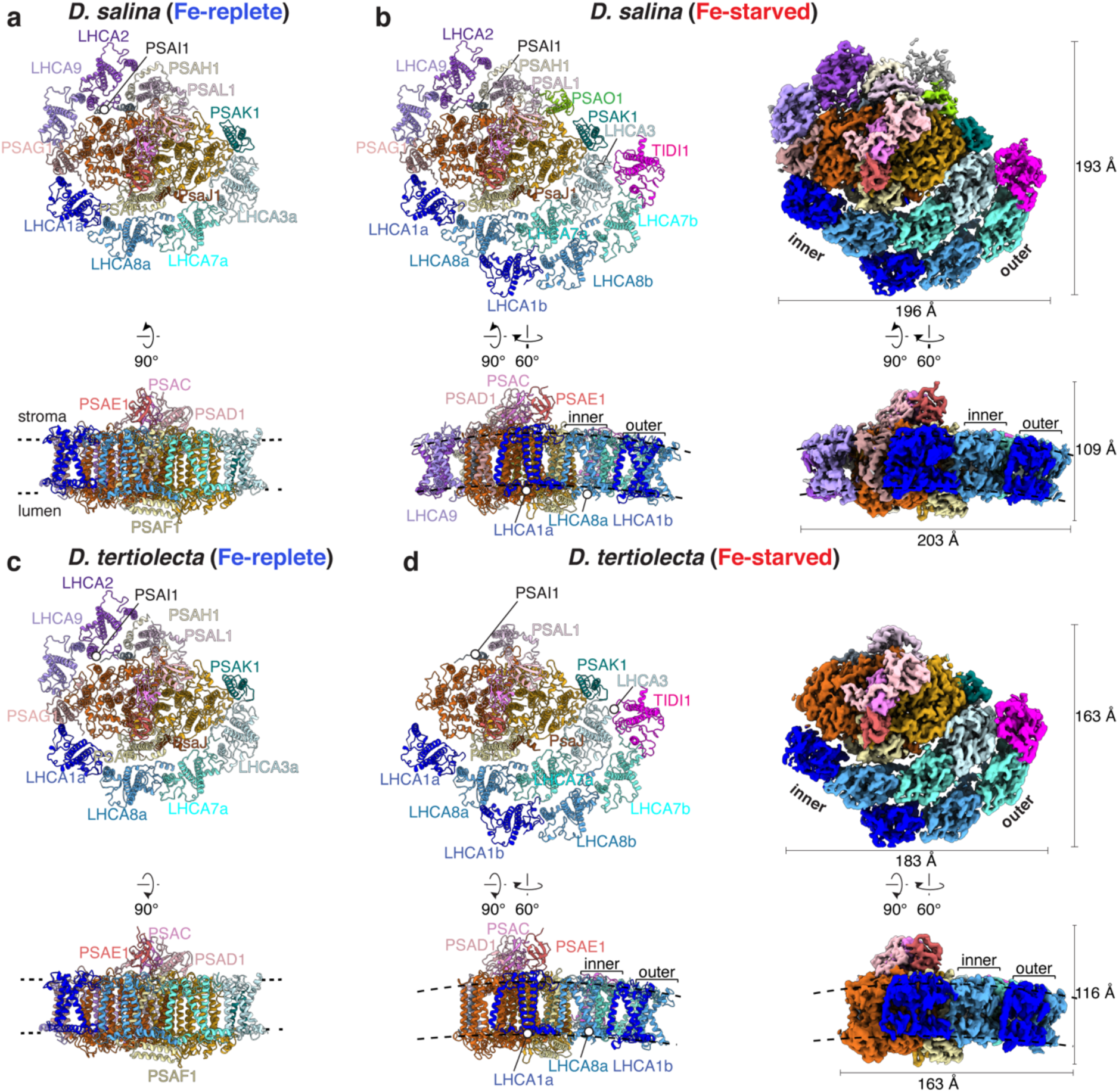
Comparison of the Fe-replete and Fe-starved *Dunaliella* PSI-LHCI supercomplexes. Stromal view (top) or side view (bottom) of the PSI-LHCI supercomplexes from *D. salina* (**a**,**b**) and *D. tertiolecta* (**c**,**d**). (**a**) Fe-replete *Ds*PSI-LHCI_1_ supercomplex and (**c**) Fe-replete *Dt*PSI-LHCI_1_ supercomplex in ribbon mode. **b**,**d,** Fe-starved *Ds*PSI-LHCI_2_ supercomplex (**b**) and *Dt*PSI-LHCI_2_ supercomplex (**d**) in ribbon mode (left) and the composite cryo-EM density map (right). Subunits are shown in different colors and labeled. Membrane-extrinsic subunits PSAC, PSAD and PSAE are labeled only in the side view. Black dashed lines indicated the membrane-spanning region of the PSI-LHCI supercomplexes.

LHCA2 is unique among the LHCI subunits in that it contains a fourth transmembrane helix whose C terminus is exposed to the stroma. We modeled several additional amino acids at the C-terminal region of LHCA2 in the *Ds*PSI-LHCI_1_ supercomplex not previously reported, 18 additional residues relative to the *D. salina* PSI-LHCI supercomplex (PDB: 6SL5) and 8 to 19 additional residues compared to PSI structures from other green algae (Extended Data Fig. 7, (*26*, *28–31*)). The extended stromal region interacts with the PSAI1 subunit through hydrophobic side chains, ascribing a function to PSAI1 in attachment and stability of the LHCA2-LHCA9 dimer (Extended Data Fig. 7).

We identified densities for 171 chlorophyll (Chl) *a*, 1 Chl *a*’, 13 Chl *b*, 46 carotenoids (Cars), 3 Fe_4_S_4_, 2 phylloquinones, and 56 bound lipid molecules in the *Ds*PSI-LHCI_1_ supercomplex (Extended Data Fig. 8, 9, Extended Table 2). Although *D. tertiolecta* and *D. salina* are separated by ∼250 million years of evolution and inhabit distinct ecological niches, coastal Arctic vs. hypersaline desert, the PSI-LHCI supercomplexes from the Fe-replete condition are highly similar, underscoring the high conservation of PSI subunit organization and antenna arrangement in the *Dunaliella* radiation (Fig. 2). For *D. tertiolecta*, we identified 167 Chl *a*, 1 Chl *a*’, 13 Chl *b*, 42 Car, 3 Fe_4_S_4_, 2 phylloquinones and 42 lipid molecules (Extended Data Fig. 8, 10, Extended Table 2).

### An additional, TIDI1-containing, LHCI tetramer in Fe-starved Dunaliella PSI-LHCI supercomplexes

Cryo-EM analysis of the PSI-LHCI supercomplexes from Fe-starved cells revealed major populations (∼60% and 30% for *D. salina* and *D. tertiolecta,* respectively) distinct from the PSI- LHCI_1_ structures found in the Fe-replete cells. The resulting structures (hereafter named PSI- LHCI_2_) reached overall resolutions of 2.9 Å for *D. salina* and 2.7 Å for *D. tertiolecta*. The PSI- LHCI_2_ supercomplexes have an additional crescent-shaped LHCI tetramer with associated pigments that is evident in both *Dunaliella* spp. grown in Fe-starved conditions (Fig. 2, b and d, Extended Data Fig. 11, 12, 13, and 14, Extended Table 3 and 4). Both *Dunaliella* spp. PSI-LHCI_2_ supercomplexes contain 10 PSA subunits, an inner LHCI tetramer arranged as in the PSI-LHCI_1_ supercomplex structure, and an outer LHCI tetramer (Fig. 2, b and d, Extended Table 3 and 4).

Like the PSI-LHCI_1_ supercomplexes, these PSI-LHCI_2_ supercomplex densities are more rigid around the PSI core and more flexible in the crescent-shaped LHCI tetramer regions, with the outer LHCI tetramer being even more flexible (Fig. 2, b and d, Extended Data Fig. 11, 12, and 13). Local refinement leading to a 3.3 Å global resolution for this region allowed us to identify LHCA1, LHCA8 and LHCA7, with the fourth subunit identified as TIDI1 after even further local refinement (Extended Data Fig. 15 and 16). The larger size of the *Ds*PSI-LHCI_2_ supercomplex versus the *Dt*PSI-LHCI_2_ supercomplex in the biochemical preparations can be explained, in part, by the presence of PSAG1, PSAH1, PSAO1, and the LHCA2-LHCA9 dimer in the *Ds*PSI-LHCI_2_ supercomplex, but not the *Dt*PSI-LHCI_2_ supercomplex (Extended Data Fig. 3). The biochemical preparations from Fe-starved *D. tertiolecta* cells contained a minor population of PSI-LHCI_1_ supercomplexes (∼20%, resolved to 3.1 Å) containing PSAG1, LHCA2 and LHCA9 (Extended Data Fig. 12), indicating that the absence of these subunits in the *Dt*PSI-LHCI_2_ complex is not trivially explained by dissociation during sample preparation or vitrification.

In terms of pigments and cofactors, the *D. salina* and *D. tertiolecta* PSI-LHCI_2_ supercomplexes contain, respectively, 218 and 184 Chl *a*, 23 and 20 Chl *b*, 59 and 47 Car, 56 and 26 lipids, plus 3 Fe_4_S_4_, 1 Chl *a*’, and 2 phylloquinones in the reaction center (Extended Data Fig. 8, 9, 10 and Extended Table 3 and 4). The majority of the pigment differences between *D. salina* and *D. tertiolecta* correspond to pigments associated with the three extra PSA and two extra LHCA subunits in the *Ds*PSI-LHCI_2_ supercomplex (Extended Table 3 and 4).

Two crescent-shaped tetramers are a common feature in chlorophyte PSI-LHCI supercomplexes, as observed in *Chla. reinhardtii*, *Bryopsis corticulans*, and *Chlorella ohadii* (*28*, *29*, *32*). But the outer LHCI tetramer in *Dunaliella* spp. is distinct. First, a conventional LHCA protein is replaced by TIDI1, and second, its orientation within the supercomplex is unique (Fig. 2, b and d). The inner LHCI tetramer, containing LHCA1, LHCA8, LHCA7 and LHCA3, has a similar association with the PSI core in all reported PSI structures in the Viridiplantae clade (Extended Data Fig. 17, a and b) (*22*, *33*). The outer LHCI tetramer of the *Dunaliella* spp. PSI- LHCI_2_ supercomplexes is rotated towards PSAK1 by ∼33 degrees, whereas in other reported green algal structures, the outer LHCI tetramer is aligned with the inner LHCI tetramer (*28*, *29*, *32*) (Extended Data Fig. 17a). Outer LHCI tetramer rotation occurs also in a bryophyte (*Physcomitrium patens*) large PSI-LHCI supercomplex, but through a very distinct mechanism that involves association with additional LHCII oligomers (Extended Data Fig. 17b) (*31*, *34*). In contrast, in *Dunaliella* spp., LHCA1 of the outer LHCI tetramer (hereafter LHCA1b) interacts with inner LHCI tetramer subunits (see below) that are closer to the proximal side (Fig. 2 and Extended Data Fig. 17b). These distinct arrangements of the outer tetrameric LHCA sub-complexes in divergent organisms may represent independent convergent mechanisms for adjusting PSI accessory light-harvesting antenna.

The *Dunaliella* outer LHCI tetramer forms a crescent-shaped arrangement similar to the inner LHCI tetramer. Superimposition of the inner and outer LHCI tetramers results in a root mean square deviation (RMSD) of 0.36 and 0.66 Å (over 223 and 224 aligned core Cα atoms) for *Ds*PSI- LHCI_2_ and *Dt*PSI-LHCI_2_ supercomplexes, respectively (Extended Data Fig. 18a). This suggests that the relative positions of each of the LHCA monomers are well conserved within the inner and outer LHCI tetramers. Interestingly, when the outer LHCI tetramers of *Ds*PSI-LHCI_2_ and *Dt*PSI- LHCI_2_ supercomplexes are superimposed, the RMSD is 0.56 Å (over 223 aligned core Cα atoms) (Extended Data Fig. 18a). This indicates the high conservation of the arrangement of LHCA proteins in the outer LHCI tetramer between two divergent *Dunaliella* spp. in Fe starvation. However, the outer LHCI tetramer of the PSI-LHCI_2_ supercomplexes are slightly less curved than the inner tetramer in both *Dunaliella* spp. (Extended Data Fig. 18b), which are similarly observed in other PSI-LHCI supercomplexes with a double LHCI tetramer antenna arrangement (*28*, *31*).

The inter-subunit interactions in the inner and outer tetramer are also conserved in the two *Dunaliella* spp. (Fig. 3, Extended Data Fig. 19). On the proximal side, LHCA1b interacts with both LHCA8 and LHCA7 in the inner LHCA tetramer (hereafter LHCA8a and 7a) (Fig. 3, Extended Data Fig. 19, a and e). The lumen side of helix C in LHCA1b interacts with the BC loop of LHCA8a through hydrophobic side chains, while its stromal side interacts with the AC loop of LHCA8a through charged side chains. The latter interaction is only evident in the *Ds*PSI-LHCI_2_ supercomplex, but is poorly resolved in the *Dt*PSI-LHCI_2_ supercomplex (Fig. 3, Extended Data Fig. 19, a and e). The N-terminus of LHCA1b uses hydrophobic side chains to interact with the AC loop of LHCA7a on the stromal side (Fig. 3, Extended Data Fig. 19, b and f). At the distal side of the inner crescent, LHCA7 of the outer LHCI tetramer (hereafter named LHCA7b) interacts with LHCA3 through both polar and non-polar side chains at its stromal side N-terminus (Fig. 3, Extended Data Fig. 19, c and g). In the *Ds*PSI-LHCI_2_ structure, we note an interaction between the lumen side C-terminus of LHCA7b and the AC loop of LHCA3, but this is not well-resolved in our EM density of the *Dt*PSI-LHCI_2_ structure (Fig. 3, Extended Data Fig. 19, c and g). Several pigments, including Chl and Car, and non-protein densities attributed to lipids were observed in these interfaces, suggesting their contribution to the attachment of the outer and inner LHCI tetramers (Extended Data Fig. 8).

**Fig. 3.**
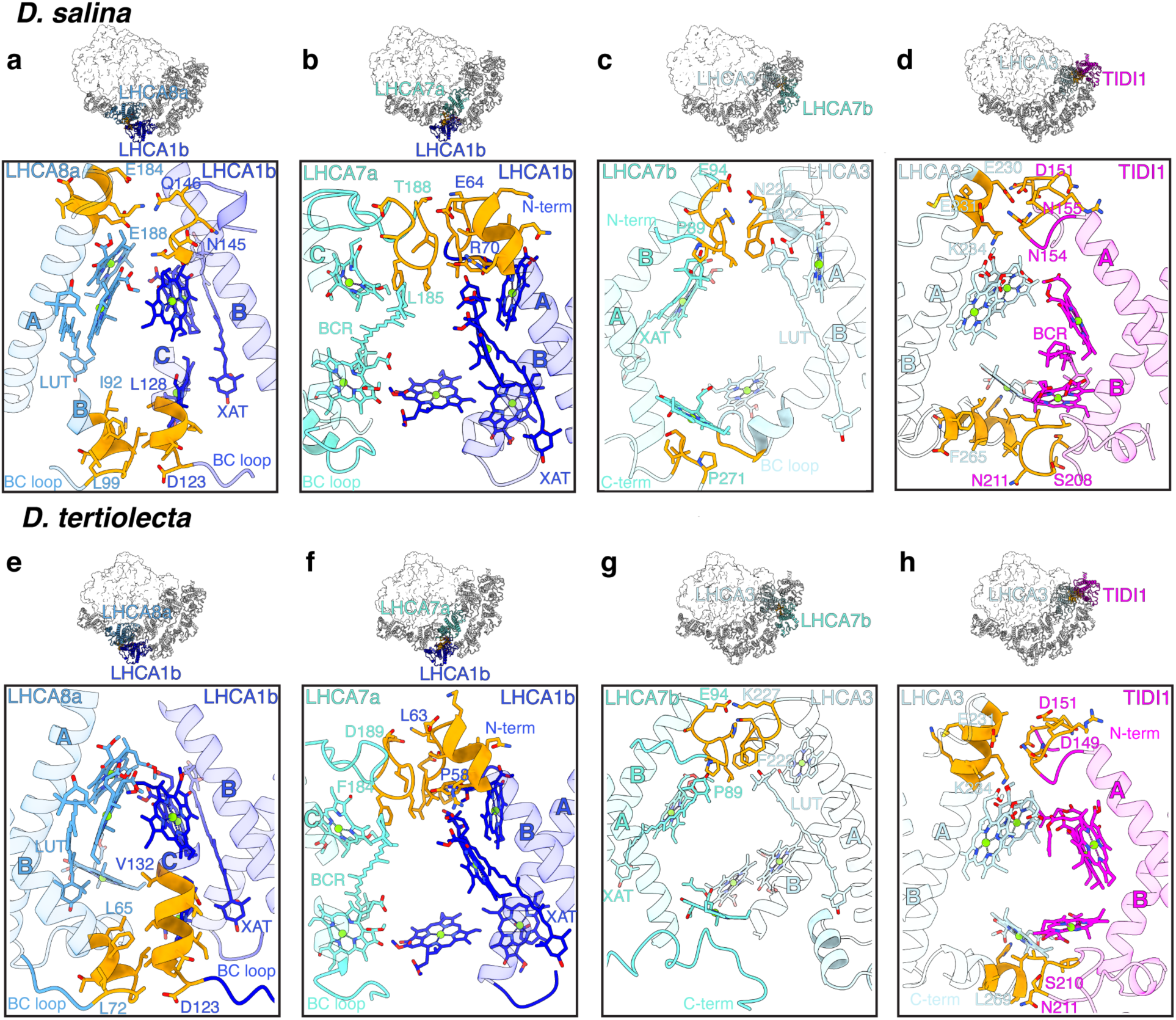
The interactions between the inner LHCI tetramer and the outer LHCI tetramer in the *Ds*PSI-LHCI_2_ and *Dt*PSI-LHCI_2_ supercomplexes. **(a-h)** Top: The interaction regions between the inner and outer LHCI tetramers in the overall *Ds*PSI-LHCI_2_ (**a-d**) and *Dt*PSI-LHCI_2_ (**e-h**) supercomplexes. Boxed areas are enlarged in the indicated panels. Bottom: protein-protein, pigment-pigment, and protein-pigment interactions between: LHCA8a and LHCA1b (**a**,**e**), LHCA7a and LHCA1b (**b**,**f**), LHCA3 and LHCA7b (**c**,**g**), LHCA3 and TIDI1 (**d**,**h**). The residues involved in the interactions are highlighted as yellow stick representations.

### TIDI1 is a key constituent of the second LHCI tetramer

We confirmed the location of TIDI1 as the most distal subunit in the outer LHCI tetramer of the PSI-LHCI_1_ supercomplex using TIDI1-specific features relative to LHCA3 and other LHCAs. These TIDI1-specific features include an extended BC loop with the diagnostic PFXGX_2_PF motif as well as primary sequence differences (Fig 2, Extended Data Fig. 20, b and d, Extended Data Fig. 15 and 16) (*13*, *35*). However, the extended proline-rich region between L31 and S147 for *Ds*TIDI1 and L129 for *Dt*TIDI1 (*13*) was too flexible and could not be resolved. The presence of TIDI1 in the outer LHCI tetramer instead of LHCA3 is consistent with the reduced abundance of LHCA3 relative to LHCA1, LHCA7 and LHCA8 in whole cell proteomics analysis (Fig. 1, e and f). Like other members of the LHC family, TIDI1 has three transmembrane helices (Fig. 4a, Extended Data Fig. 20a), and its interactions via its stromal AC loop with the N-terminus of the adjacent LHCA7b subunit are similar to the LHCA3-LHCA7a interaction (Fig. 4b, Extended Data Fig. 21b). The TIDI1-specific BC loop contacts helix C of LHCA3 at the lumenal side in both *Dunaliella* spp. PSI-LHCI_2_ supercomplexes, with polar side chains contributing to the interaction (Fig. 3, d and h, Extended Data Fig. 21). Charged side chains are also involved in the stromal side interaction between the N-terminus of TIDI1 and the AC loop of LHCA3, reminiscent of the interaction of the N-terminus of LHCA3 with the PsaA subunit for association of the inner-tetramer with the PSI core (Fig. 3, d and h, Extended Data Fig. 21). The stroma- and lumen-side TIDI1-LHCA3 interactions suggests a role for TIDI1 in positioning the outer LHCI tetramer with respect to the inner LHCI tetramer.

**Fig. 4.**
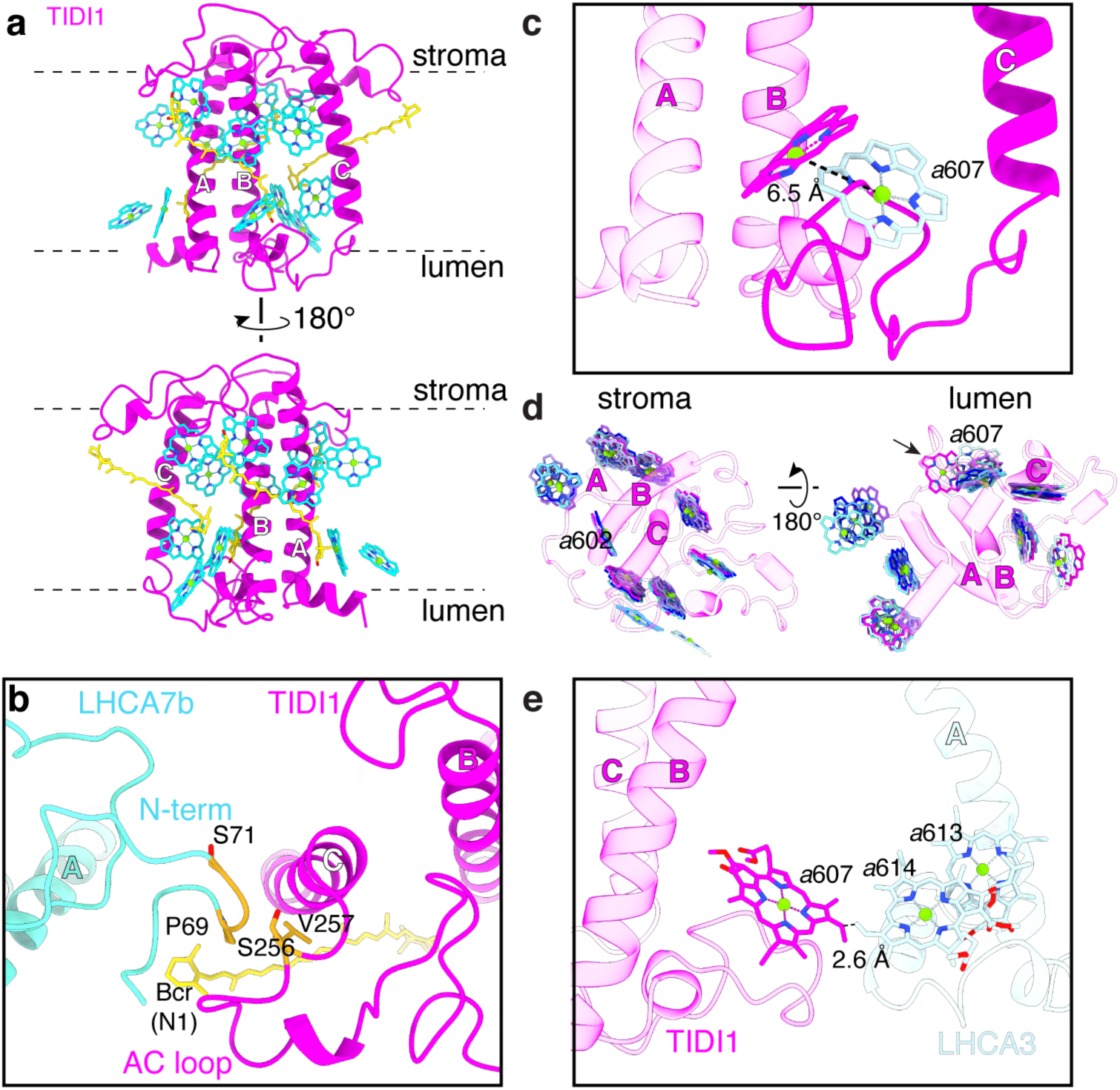
The extended BC loop of *Ds*TIDI1 differently positions Chl 607. **a**, Cartoon representation of *Ds*TIDI1. Chls are shown as cyan stick representation, with the central Mg atoms shown as spheres and numbered according to the conserved sites in spinach LHCII (PDB code 1RWT). Carotenoids are shown as yellow-colored stick representation. **b**, Interactions between *Ds*TIDI1 and *Ds*LHCA7b in the outer LHCI tetramer are colored in orange. Residues involved in the interaction are shown as orange sticks. **c**, Comparison of Chl *a*607_TIDI1_ position against position of *a*607_LHCA3_ from *D. salina* from the lumenal side. Distance between molecules is shown as dashed lines. **d**, Comparison of *Ds*TIDI1 Chl positions against Chl positions of *Ds*LHCA1, *Ds*LHCA2, *Ds*LHCA3, *Ds*LHCA7, *Ds*LHCA8, and *Ds*LHCA9 from *D. salina* on the stromal (left) and lumenal side (right). **e**, Side view of *D. salina a*607_TIDI1_ (magenta) and its distance shown as dashed lines to *a*614_LHCA3_ (light blue).

We modeled three Cars in TIDI1 in the same location as those in LHCA3 in both structures, and 13 Chls in *D. salina* and 12 in *D. tertiolecta* (Fig. 4a, Extended Data Fig. 20a). As in other LHC subunits, the Chl-Car interactions occur primarily at the stromal pigment layer, contributing to the asymmetric enrichment of pigments at the stromal side. All Chl positions of TIDI1 were equivalent to those in LHCA3, except for Chl *a*607 (hereafter *a*607_TIDI1_) on the lumenal side. Compared to *a*607_LHCA3_, *a*607_TIDI1_ is shifted 6.5 Å or 8.2 Å (Mg to Mg distance) and rotated by ∼ 67 or 61 degrees in the *D. salina* and *D. tertiolecta* supercomplexes, respectively (Fig. 4c, Extended Data Fig. 20c). The position and orientation of *a*607_TIDI1_ are unique to TIDI1 and have not been observed in any other LHC protein (Fig. 4d, Extended Data Fig. 20d and Fig. 22). *a*607_TIDI1_ is at the interface of TIDI1 and LHCA3 at the lumen side and is 2.6 Å and 1.8 Å (in *D. salina* and *D. tertiolecta*, respectively) from *a*614_LHCA3_ (Fig. 4e, Extended Data Fig. 20e), close enough for energy transfer between TIDI1 and LHCA3. The similar positions and orientations of the majority of the pigments in TIDI1 to LHCA3 supports its evolution from LHCA3 and functional equivalency in energy transfer.

### Multiple evolutionary paths to an outer LHCI tetramer

The double LHCI tetramer arrangement occurs in two different ways (aligned versus severely rotated) in the Viridiplantae clade. In many chlorophyte algae, the expansion of the *LHCA* genes gave rise to LHCA4, LHCA5, and LHCA6 (*36*). LHCA5 and LHCA6 are specialized with extended lumenal C-terminal regions that facilitate the association of the outer and inner LHCI tetramers to form the parallel outer LHCI tetramer arrangement (*28*, *29*, *32*) (Extended Data Fig. 17c). In *P. patens*, the subunits of the inner LHCI tetramer are re-used in the outer LHCI tetramer to form a very tilted offset outer LHCI tetramer that connects also with LHCB9 and an LHCII trimer on the opposite side (*31*) (Extended Data Fig. 17). Here, we identify a third arrangement in *Dunaliella* spp. where the outer LHCI tetramer is facultative. It is induced in Fe starvation by selective stoichiometric adjustment of three inner tetramer subunits (LHCA1, LHCA8 and LHCA7) along with the innovation of a unique LHCA-type protein (TIDI1) (Fig. 2, b and d), likely evolved from an ancestral LHCA3 (*13*). The result is a novel LHCI tetramer whose orientation is also tilted and offset, albeit less so than in bryophytes (*31*). These comparisons underscore the relevance of the extended C-termini in the chlorophyte LHCA5 and LHCA6 for the parallel tetramer arrangements.

## Discussion

### Convergent evolution of modified PSI antenna in Fe-starved algae

Poor Fe bioavailability impacts primary productivity at a global scale (*1*, *2*, *37*) because of loss of the photosynthetic apparatus. At a biochemical level, the prime targets are PSI and ferredoxin because of their abundance and high Fe content, and their reduced abundance is a classic biomarker for the Fe-starved state (*4*, *6*, *7*, *9*, *38–40*). A well-known adaptation, occurring in both cyanobacteria and some eukaryotic algae, is the replacement of iron-containing ferredoxin by iron-independent flavin-containing flavodoxin, which reduces the Fe quota of the cell, yet enables continued photosynthesis-driven reductive metabolism (*13*, *21*, *40–43*). Fld is reduced by electrons from PSI whose abundance and hence throughput is impacted by Fe starvation induced PSI retrenchment. In cyanobacteria, the *isiB* gene encoding Fld is co-expressed with *isiA*, which encodes a Chl-binding protein (also named CP43’ because of its sequence relationship to CP43 (PsbC) in PSII) (*21*, *44–46*). Increased expression of *isiA* in Fe-starved cyanobacteria promotes remarkable reorganization and expansion of the PSI-antenna complex to compensate for the retrenchment of the PSI core complex (*47*, *48*).

The co-expression of IsiA with IsiB is likely required to maximize anabolic metabolism in the Fe-starved cyanobacteria. A survey of 159 cyanobacterial genomes showed that 76 do contain the *isiA* gene and of those, the vast majority (71) also encode a Fld (Extended Data Fig. 23, Extended Dataset 3). In a majority (56) of the genomes that contain both, the *isiA* and *isiB* genes are adjacent and colinear, possibly in an *isiAB* operon as is the case in *Synechococcus elongatus* (PCC 7942) (*49*). Fld is not present in streptophytes, but does occur in some green algae, including *Dunaliella* spp., and typically it occurs with a TIDI1-homologue (*13*, *50*) (Fig. 1b). The co-occurrence of Fld and TIDI1 in diverse green algae (separated by ∼650 million years (*13*)) evokes an adaptive strategy reminiscent of the cyanobacterial IsiA/IsiB (Fld) system. Introduction of Fld into crop plants as a strategy for increasing photosynthetic resilience has been discussed, but thus far it has not yielded significant improvements in low Fe situations (*51*). PSI antenna reorganization may be one approach to achieving this goal.

The metabolic adaptations in play in eukaryotes have not been well-investigated beyond documentation of PSI structural and functional reduction (*8*, *52*, *53*). Here we fill this gap by structural analysis of PSI-LHCI supercomplexes from >250 million years divergent *Dunaliella* spp. from distinct saline environments, Arctic ice (*D. tertiolecta*) and desert hypersaline lake (*D. salina*) that exhibit unusual productivity with low Fe supplementation (Extended Data Fig. 1, (*13*)). We find that increased expression of TIDI1 under Fe starvation is correlated with the formation of a novel TIDI1-containing LHCI tetramer that associates with the PSI-LHCI supercomplex to effect a larger antenna analogous to the long known rearrangement of the prokaryotic PSI antenna (*12*, *13*, *47*, *48*) (summarized in Fig. 5).

**Fig. 5.**
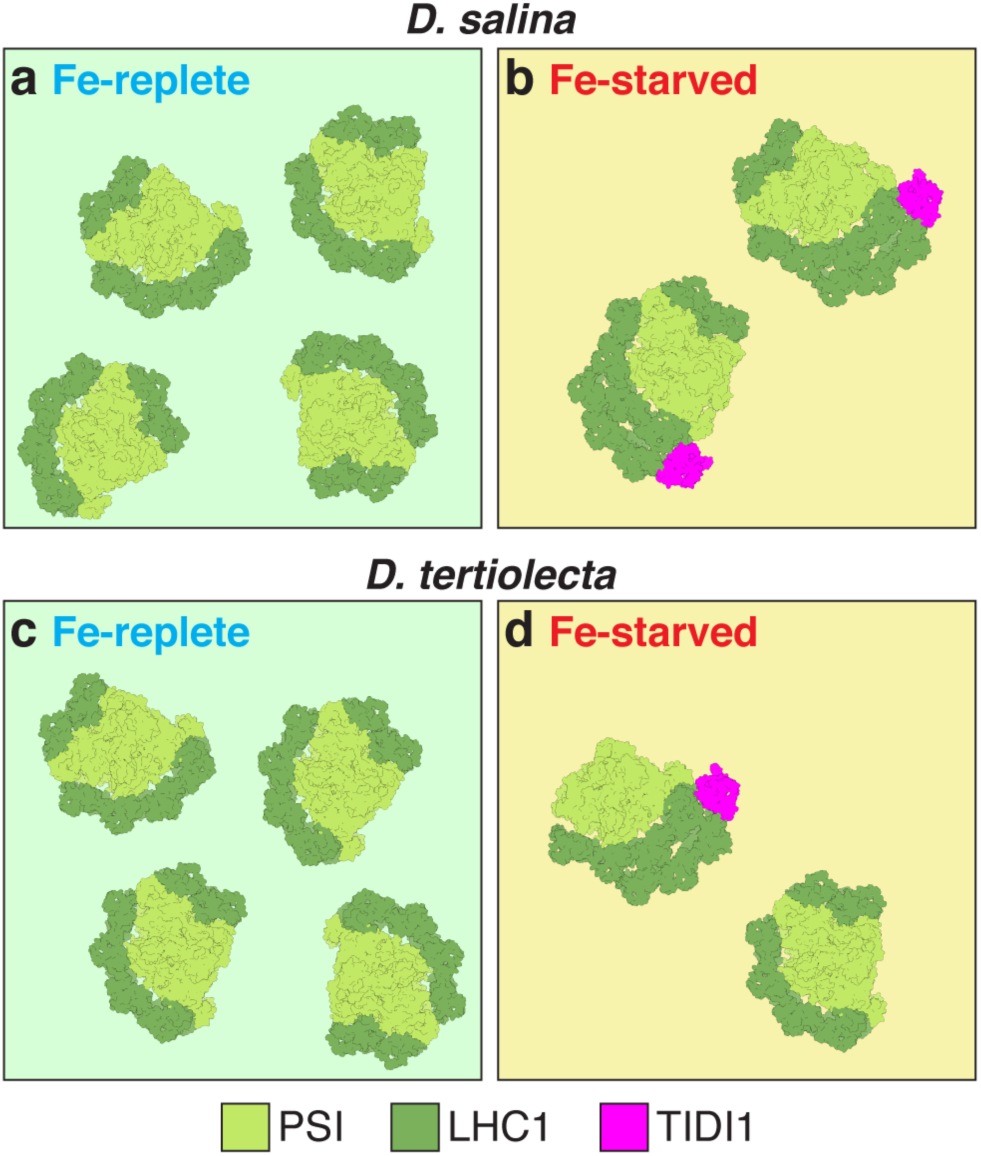
Fe-nutrition responsive modification of PSI-LHCI antenna in *Dunaliella* spp. **a**,**c**, PSI-LHCI supercomplexes in Fe-replete cells have a single LHCI tetramer (dark green) composed of four subunits in *D. salina* (**a**) and *D. tertiolecta* (**c**). When cells become Fe-starved, the PSI-LHCI supercomplexes has an additional LHCI tetramer (dark green) also composed of four subunits with the novel, Fe starvation induced chlorophyll-binding protein, TIDI1 (pink), in *D. salina* (**b**) and *D. tertiolecta* (**d**). The PSI core subunits are colored in light green. The *Dt*PSI-LHCI_2_ does not include the PSAO1, PSAG1, PSAH1, LHCA2, and LHCA9 subunits when compared to those from *D. salina* cells. Fe-starved *D. tertiolecta* samples include a small fraction of *Dt*PSI-LHCI_1_ supercomplexes, as shown.

### Fe starvation induced antenna diversity in other eukaryotic algae

The unprecedented structural re-arrangement of the LHCI antenna in Fe-starved *Dunaliella* spp. was foreshadowed by whole-cell proteomic analyses (Fig. 1, d and e), which revealed uniform heterogeneity in both species in the response of three of the six LHCA subunits during the Fe starvation-dependent retrenchment of the photosynthetic apparatus. LHCA1, LHCA7 and LHCA8, which are constituents of the novel outer LHCI tetramer, were retained, but LHCA2 and LHCA9, which make up the dimer and LHCA3 that connects the tetramer to the PSI core, decreased in proportion to the decrease in PSI core subunits. The structures of the PSI- LHCI_2_ supercomplex explain the discrepancy because three of the four subunits of the inner tetramer are reused in the new TIDI1-containing tetramer (Fig. 2, b and d). Whole-cell proteomics is a valuable tool for assessing the physiological relevance of unusual subunit compositions in isolated complexes. Although we did not undertake absolute quantification of each of the subunits in the Fe-replete cells, we note that PSAG1, PSAL1, PSAH1, PSAI1, LHCA2 and LHCA9 were captured in the proteomics dataset of both species (Fig. 1, e and f, Extended Dataset 2), and indeed these subunits were found in both Fe-replete structures (Fig. 2, a and c). We did not observe particles corresponding to the minimal PSI-LHCI supercomplex found in other work with *D. salina* (*27*), and we wonder whether subunit dissociation may occasionally be a problem during isolation or vitrification. Interestingly, most of the same subunits (PSAG1, PSAH1, LHCA2 and LHCA9) are absent from the *Dt*PSI-LHCI_2_ supercomplex in our present work as well. Nevertheless, we found PSI-LHCI_1_ supercomplexes in the Fe-starved *D. tertiolecta* PSI-LHCI preparations (Fig. 2d, Extended Data Fig. 12) with the typical PSA and LHCA subunit compositions, including PSAG1, PSAH1, LHCA2 and LHCA9. This result suggests that the absence of these subunits in the *Dt*PSI-LHCI_2_ supercomplex is likely not an isolation or vitrification artefact. The absence may reflect genuine in vivo heterogeneity, which is not evident in the whole cell proteomics because of the substantial amount of the *Dt*PSI-LHCI_1_ species.

The differential response of a subset of the LHCA proteins in *Dunaliella* spp. was distinct from observations of the well-studied chlorophyte *Chla. reinhardtii,* where all the subunits of the supercomplex are proportionally decreased under Fe-limited growth (Extended Data Fig. 24b) (*54*, *55*). *Chla. reinhardtii* and *Chro. zofingiensis* have nine distinct and orthologous LHCA subunits versus six LHCA subunits in *Dunaliella* spp. (*13*, *21*, *56*). Although the structure of the *Cz*PSI-LHCI supercomplex has not been determined, it is likely to be similar to that of the other chlorophyte algae (discussed above) with two aligned tetramers, an inner one made up of LHCA1, LHCA8, LHCA7 and LHCA3 and an outer one made up of LHCA1, LHCA4, LHCA5 and LHCA6 (*28*, *29*, *32*). Whole-cell proteomics of *Chro. zofingiensis*, whose genome also encodes both FLD1 and TIDI1 (which are coordinately upregulated in Fe-deficiency) (Extended Data Fig. 24c), also revealed selective differences in retention of some LHCA proteins (LHCA1, LHCA8, LHCA7, LHCA4, LHCA5, LHCA6) and proportional (to PSA subunits) losses of others (Extended Data Fig. 24) (*57*). We anticipate that structural analysis of the Fe-starved PSI-LHCI supercomplex from *Chro. zofingiensis* will reveal another novel quaternary arrangement of LHCI tetramers. In the diatom *Phaeodactylum tricornutum*, which encodes an FLD1 ortholog, the mRNA abundances for four Chl-binding proteins are upregulated in Fe-limitation, suggesting that there may be remodeling of a diatom-specific PSI-LHCI supercomplex under Fe starvation (*39*, *40*).

## Assembly of a second LHCI tetramer

For both *D. tertiolecta* and *D. salina*, the arrangement of the PSI core and the inner LHCI tetramer subunits in the PSI-LHC_2_ supercomplexes is almost identical to that in the PSI-LHCI_1_ supercomplexes (Fig. 2, Extended Data Fig. 18). This observation is compatible with a model in which the PSI-LHC_2_ supercomplex is assembled by the addition of the new tetramer to the pre-existing PSI-LHCI_1_ supercomplexes. De novo synthesis of the entire supercomplex is unlikely, given that *PSA* transcript abundances are decreased under Fe starvation (*13*). The replacement of LHCA3 with TIDI1 may determine the position of the new tetramer on the outside rather than in direct association with the PSI core. Although we made a substantial effort to resolve the extended proline-rich region of TIDI1, including the use of a GraFix strategy for Fe-starved *D. tertiolecta*, we cannot assess its role in determining subunit-subunit interactions that promote assembly. The interaction of LHCA3 with the PSI core complex may be a common target for post-translational events affecting PSI-LHCI function. In *Chla. reinhardtii*, the N-terminus of LHCA3 is removed in low Fe, perhaps initiating the subsequent degradation of LHCA3 (*58*).

Together with the introduction of flavodoxin (*59*), eukaryotic LHCI antenna remodeling presents a new engineering opportunity for improving PSI performance under low Fe nutrition stress, and in so doing can improve crop and oceanic productivity in environments that suffer from chronic Fe limitation.

## Materials and Methods

### Strains and Culture Conditions

*Dunaliella tertiolecta* and *Dunaliella salina* were provided by Uri Pick. The medium composition and growth conditions were modified from (*60*). The NaCl concentration was reduced to 0.5 M, and the solution was de-metalated by passage through a Chelex 100 Resin (BioRad) column prepared according to (*61*). Fe was added to a final concentration of 15 or 0.15 µM FeCl_3_ with 6 µM EDTA, for the Fe-replete or the Fe-depleted conditions, respectively. All glassware was washed in 6 N HCl and rinsed several times with Milli-Q water. Cultures (100 to 500 mL) were grown in 250 mL to 1 L Fernbach flasks on shaking platforms (140 rpm) at 24°C. Continuous light was provided by cool white fluorescent bulbs (4,100 K) and warm white fluorescent bulbs (3,000 K) in a 2:1 ratio at 100 µE m^-2^ s^-1^. Cells were counted with a Beckman Coulter Multisizer 3 with a 50-µM orifice (Beckman Coulter).

### Elemental analysis

Elemental analysis was determined by ICP-MS/MS (*62*) with minor modifications. *Dunaliella* cells (6 x 10^6^ cells/mL) from logarithmic growth (between 1 and 2 x 10^6^ cells/mL) were collected by centrifugation at 1,680 x*g* for 3 min in a 50-mL Falcon tube. Cells were washed twice in a solution containing 1 mM Na_2_-EDTA, pH 8, 0.5 M Chelex treated NaCl and twice in Chelex-treated 0.5 M NaCl and collected by centrifugation. The cell pellet was digested with 143 µL of 70% nitric acid (Optima grade, Fisher, A467-500) at 65°C for 2-4 hours and diluted to a final nitric acid concentration of 2% (v/v) with Milli-Q water. Elemental analysis was by ICP-MS/MS on an Agilent 8900 Triple Quadrupole instrument, in comparison with an environmental calibration standard (Agilent 5183-4688), a sulfur (Inorganic Ventures CGS1) and a phosphorus standard (Inorganic Ventures CGP1). ^89^Y (Inorganic Ventures MSY-100PPM) was used as an internal standard. The levels of all analytes were determined in MS/MS mode.

### Chl content

Chl content was collected from 1 mL of culture (1-2 x 10^6^ cells/mL) by centrifugation at 21,130 x*g* for 1 min at 4°C and extracted with 80:20 (v/v) acetone to water. Debris was removed centrifugation at 21,130 x*g* for 5 min at 25°C and the absorbance of the supernatant was measured at 647 nm and 664 nm to calculate Chl according to the extinction-coefficients of (*63*). For isolated membranes, Chl was extracted from 5 µL of the preparation (see below).

### Preparation of TIDI1 antiserum

Affinity-purified rabbit polyclonal antibodies specific for TIDI1 were raised at Labcorp (Madison, Wisconsin, USA) against the peptide NH_2_-PEPKKGSAFKGY-COOH (*12*) according to their protocols.

### Immunodetection

10 mL of cultures at a density of 1-2 x 10^6^ cells/mL was collected by centrifugation at 1,680 x*g* at 4°C and stored –80°C. Cell pellets were resuspended in 300 µL 10 mM sodium phosphate, pH 7.0. Cells were broken by freeze-thaw cycling according to (*64*). Protein concentrations were determined by Pierce BCA assay against a bovine serum albumin standard (Thermo Fisher Scientific). Proteins were separated by SDS-PAGE and immunodetection as described by (*64*) except a different transfer buffer (25 mM Tris, 192 mM glycine, 20% (v/v) methanol) was used and TBS (10 mM Tris-HCl, 150 mM NaCl, pH 7.5) with 0.05% (w/v) Tween 20 (TBS-T) was used to dilute the antibodies and wash the membranes. Primary antibody dilutions were as follows: CF_1_ 1:50,000 (*65*), FDX1 1:500 (*66*), FLV1 1:500 (*13*), PSAF1 1:1000 (*67*), TIDI1 1:1000 (this publication). For visualization of bound antibody, washed membranes were incubated in a 1:6000 dilution of goat anti-rabbit secondary antibody (Southern Biotech) conjugated to alkaline phosphatase in 3% (w/v) nonfat dried milk in 1xPBS with 0.1% (w/v) Tween 20 and visualized by incubation for 0.5–1 min in 10 mL alkaline phosphatase buffer.

### Proteomic analyses

Cells (1 x 10^8^) were collected by centrifugation at 1,680 x*g* at 4°C for 5 min and resuspended in 300 µL 10 mM phosphate, pH 7.0, frozen, and shipped for analysis at Environmental Molecular Sciences Laboratory (EMSL). The cell samples were fractionated and labelled with tandem mass tags as published in (*68*). Protein identities and abundances were quantified from all LC-MS/MS spectra using the software MSGF+ (*69*), and reporter ion abundances were extracted from each spectrum using MASIC (*70*) to measure peptide abundance. Each peptide abundance was normalized to the mean central tendency of all peptide abundances per sample. Downstream statistical analysis of three separate cultures were conducted on log_2_ abundance of each sample. The protein names for *Chla. reinhardtii* and *Dunaliella* spp. are based on the updated nomenclature from (*56*).

### Isolation of PSI-LHCI supercomplexes

*Dunaliella* Fe-starved and Fe-replete thylakoid membranes were first purified by sucrose cushion centrifugation as described previously (*34*) with modifications. Cultures containing 1-3 x 10^6^ cells/mL were collected at 1,670 x*g* (JLA-10.500 rotor, Beckman Coulter). Cells were washed once (*D. tertiolecta*) or twice (*D. salina*) in a medium containing the respective Fe concentrations. The cell pellet was osmotically lysed on ice for 30 min in 25 mM MES-NaOH (pH 6.5), 1.5 mM NaCl, 0.2 mM benzamidine, and 1 mM ε-aminocaproic acid with intermittent gentle agitation.

To isolate the PSI-LHCI supercomplex, thylakoid membranes (400–800 µg Chl at 0.5 mg Chl/mL) were solubilized with 1% (w/v) *n*-dodecyl-α-D-maltoside (α-DDM) (Anatrace) for 30 min with gentle agitation. The unsolubilized membranes were removed by centrifugation at 20,000 x*g* for 5 min at 4°C. The proteins from the solubilized membranes were separated by density gradient centrifugation containing either a stepwise sucrose gradient [0.1–1.8 M sucrose with 25 mM MES-NaOH (pH 6.5) and 0.03% (w/v) α -DDM] for negative staining or maltose gradient for cryo-EM data collection [0.1–1.4 M maltose with 25 mM MES-NaOH (pH 6.5) and 0.03% (w/v) α-DDM at 154,300 x*g* (SW 41 Ti rotor, Beckman Coulter) for 24 h at 4°C. PSI-LHCI supercomplexes were collected dropwise from the bottom of the tube before negative staining or single-particle cryo-EM analysis.

To stabilize the complexes further including TIDI1, mild crosslinking using the GraFix procedure (*35*) was performed for a second round of cryo-EM data collection for Fe-starved *D. tertiolecta*. For GraFix, a continuous linear gradient was prepared by mixing the light (0.1 M maltose with 25 mM MES-NaOH (pH 6.5) and 0.03% (w/v) α-DDM) and heavy solutions (1.4 M maltose with 25 mM MES-NaOH (pH 6.5), 0.03% (w/v) α-DDM, and 0.075% glutaraldehyde (v/v)). The proteins from the solubilized thylakoid membranes were both separated and gently crosslinked with the GraFix technique, and the centrifugation was carried out as described earlier at 154,300 x*g* (SW 41 Ti rotor, Beckman Coulter) for 24 h at 4°C. The crosslinking reaction in the GraFix purified supercomplexes was quenched with 100 mM MES-Tris (pH 6.5).

### Fluorescence emission spectroscopy at 77K

Fractionated membranes or cultured cells at the mid-log phase were placed in a glass tube and frozen in liquid N_2_ after adjusting the Chl concentrations to 5 µg/mL. Fluorescence emission was recorded at 77 K using FluoroMax-4 spectrophotometer (Horiba Scientific) according to the previous protocol (*71*).

### Negative staining electron microscopy

The sample quality of the isolated PSI-LHCI supercomplexes was assessed through negative staining electron microscopy. Briefly, 4 µL of PSI-LHCI supercomplex samples, diluted in 25 mM MES-NaOH (pH 6.5) and 0.03% (w/v) α-DDM, was incubated for 1 min on carbon-coated Cu grids (400 mesh, Electron Microscopy Sciences) pretreated with a Tergeo-EM plasma cleaner (PIE Scientific). The grid was subsequently washed with 2% (w/v) trehalose, 0.03% (w/v) α-DDM, and in 25 mM MES-NaOH (pH 6.5) and finally stained with three 50 uL drops of 1% (w/v) uranyl formate (SPI Supplies) before subsequent blotting and drying. Negatively stained samples were imaged in a Tecnai T12 electron microscope (Thermo Fisher Scientific), operated at 120 kV, 30,000x magnification using a TemCam F-416 camera (TVIPS).

### Cryo-EM sample preparation and data collection

Cryo-EM grids were prepared with in-house fabricated carbon on gold grids, Quantifoil Au/Cu R1.2/1.3 grids 300 mesh, or Quantifoil Au/Cu R2/1 grids 300 mesh. The grids were washed with chloroform and layered with graphene oxide as previously described (*72*). Briefly, 4 µL of 1 mg/mL of polyethylenimine HCl MAX Linear MW 40k (PEI, Polysciences) in 25 mM HEPES pH 7.5 was applied and incubated on glow discharged grids for 2 min, blotted away, and washed twice with 4 µL water. The PEI-treated grids were dried with Whatman paper. 0.2 mg/ml of graphene oxide (Sigma-Aldrich, 763705) prepared in 1:5 methanol: water solution was vortexed for 10 s and precipitated at 1,000 *xg* for 1 min. 4 µL of the supernatant was applied to the PEI- treated grids for 2 min, blotted away, washed twice with 4 µL water, and dried for at least 15 min before freezing grids.

Samples were diluted with filtered 25 mM MES-NaOH (pH 6.5), 0.02% (w/v) α-DDM and 0.5% trehalose (w/v) to optimize particle distribution on the grid. Glow-discharged graphene oxide layered grids (*72*) (glow discharged for 30 s, 15 W, 64 (N/255), 10 sccm, and no purging) were mounted in a Vitrobot Mark IV (Thermo Fischer Scientific) maintained at 10°C and 100% relative humidity, with lights off. 4 µL of diluted sample was incubated for 1 min on the grid before blotting (blot time 7-9 sec and blot force 7) and immediately plunge-frozen in liquid ethane.

Frozen grids were loaded onto a Titan Krios G3i microscope (Thermo Fischer Scientific) operated at an acceleration voltage of 300 keV, equipped with a Gatan K3 summit direct electron detector operated in CDS mode, and with a GIF Quantum energy filter with a 20 eV slit width. Data collection was carried out at a nominal magnification of 81,000x in super-resolution mode (super-resolution pixel size of 0.525 Å per pixel) and with a defocus range of -0.8 um to –2.0 um. The total electron exposure was about 55-60 e/Å^2^ and each video stack comprised 50 frames. The data collection was set up using SerialEM and monitored using CryoSPARC Live (*73*). The same data collection strategy was used for all our cryo-EM datasets. 5371 movies were collected for Fe-replete *D. salina* samples, 5044 movies for Fe-replete *D. tertiolecta* samples, 6270 movies for Fe-starved *D. salina* samples, 11,299 movies for Fe-starved *D. tertiolecta*, and 11,529 movies for Fe-starved *D. tertiolecta* isolated with a GraFix strategy were collected.

### Electron microscopy data processing

All movies were aligned, gain corrected, and binned by two using MotionCorr2 implemented in RELION (v.3.0) (*74*). Contrast transfer function was computed with CTFFIND4 implemented in RELION (v.3.0). Particles were picked using crYOLO (v.1.7.7.1) with a box size of 280 pixels (*75*). Particle coordinates from crYOLO were imported into RELION3, extracted with a box size of 240 pixels (binned by two), and subjected to one round of 2D classification. Good classes that represented different views of PSI-LHCI supercomplexes were selected (326,845 particles from Fe-replete *D. salina* samples, 522,452 particles from Fe-replete *D. tertiolecta* samples, 190,745 particles from Fe-starved *D. salina* samples, and 518,929 particles from Fe-starved *D. tertiolecta*).

For the Fe-replete *D. salina* and *D. tertiolecta* datasets, particles from good 2D classes underwent a round of 3D classification using a low-pass filtered map generated from PDB 6SL5 as an initial reference and classified into three classes. The 3D class with the highest resolution was selected for iterative rounds of 3D and CTF refinement, resulting in a final global resolution of 2.8 Å for Fe-replete *D. salina* PSI-LHCI1 supercomplexes and 2.1 Å for Fe-replete *D. tertiolecta* PSI-LHCI1 supercomplexes.

Particles from good 2D classes of Fe-starved *D. salina* dataset underwent a round of 3D classification using a low-pass filtered map generated from PDB 6SL5 (*26*) as an initial reference and classified into three classes. The particles from two 3D classes that had a second LHCI tetramer were combined, resulting in 120,321 particles. These particles underwent two rounds of 3D refinement that resulted in a 3D reconstruction at final global resolution of 3.0 Å. To improve the density of the PSAO1 and TIDI1 subunits, PSAO1 and TIDI1-specific masks were used for particle subtraction. The subtracted particles underwent focused classification without alignment, resulting in a final global resolution of 3.3 Å for the PSAO1 mask and TIDI1 mask.

As for Fe-starved *D. salina* dataset, particles from 2D classes from the Fe-starved *D. tertiolecta* dataset underwent a round of 3D classification using a low-pass filtered map generated from PDB 6SL5 as an initial reference and classified into three classes. The 3D class with a second LHCI tetramer had 123,335 particles and that with a single LHCI tetramer had 108,975 particles. These particles were then subjected to iterative rounds of 3D and CTF refinement, which resulted in a final map at 2.7 Å global resolution for the PSI-LHCI_2_ supercomplex and 3.1 Å for PSI-LHCI_1_ supercomplex. To improve the density of the LHCA1 subunit, an LHCA1-specific mask was used for particle subtraction. The subtracted particles underwent focused classification without alignment resulting in a final map at 3.3 Å global resolution for the LHCA1.

The selected particles from Fe-starved *D. tertiolecta* dataset obtained using GraFix underwent 3D classification using a low-pass filtered map generated from PDB 6SL5 as an initial reference and classified into 4 classes. The 3D class with a second LHCI tetramer contained 217,073 particles. These particles then underwent iterative rounds of 3D and CTF refinements which resulted in a final map at 2.8 Å global resolution for the double LHCI tetramer structure. To improve the density for the PSAL1 and TIDI1 subunits, PSAL1 and TIDI1-specific masks were used for particle subtraction. The subtracted particles underwent focused refinement without alignment, resulting in a final map at 3.2 Å for the PSAL1 mask and TIDI1 mask.

The workflows for all the datasets are described in Figs. S4, S5, S11, S12 and S13.

### Model building and validation

The atomic models for the PSI structures were built by fitting the existing structure of *D. salina* (PDB identifier 6SL5) (*26*) as a template using PHENIX v.1.21.1 dock in map and subsequently refined with PHENIX real-space refinement (*76*). The amino acid sequences were then mutated to match the *D. salina* and *D. tertiolecta* genome and rebuilt on the basis of the cryo-EM density using COOT v.0.9.8.7 (*13*, *77*). The initial TIDI1 model was built using AlphaFold2 from the TIDI1 sequences from either *D. salina* or *D. tertiolecta*, docked into the Fe-starved *Dunaliella* PSI-LHC_2_ complexes using PHENIX dock in map, and subsequently fitted for the side chains in COOT (*78*). Models were built sequentially, starting with the maps of the refined structure, then the locally refined regions (PSAG1, PSAH1, PSAL1, PSAK1, PSAO1, LHCA1, LHCA2, and the outer tetramer), and finally the TIDI1 density. The resulting atomic models were refined using both the refined map and locally refined maps with the PHENIX real space refinement program using restraints for Chl *a* from (*79*) and the Grade Web Server (https://grade.globalphasing.org/cgi-bin/grade2_server.cgi) for other ligands. This process was followed by manual inspection to remove obvious errors. The refinement statistics are provided in Extended Table S1. Figures were prepared using UCSF ChimeraX v.1.7.1 (*80*). Interactions between subunits were analyzed through the PDBe PISA server (https://www.ebi.ac.uk/msd-srv/prot_int/cgi-bin/piserver) (*81*). The protein names for *Chla. reinhardtii* and *Dunaliella* spp. are based on the updated nomenclature from (*56*).

### isiA and fldA co-occurrence in Cyanobacteria

In order to identify the prevalence and co-occurrence of *isiA* genes encoding iron-stress-induced Chl-binding protein (CP43’) and *fldA* (or *isiB*) genes encoding flavodoxin in Cyanobacteria, a systematic survey was conducted of the annotated Cyanobacterial genomes hosted in the National Center for Biotechnology Information (NCBI) genome database (ncbi.nlm.nih.gov). All genomes and proteomes within the taxon Cyanobacteriota (blue-green bacteria) that were described in that database as “reference genomes” were downloaded for analysis (N = 219). Three genomes described only as “cyanobacterium endosymbiont” were excluded, and three non-reference genomes (*Synechococcus* sp. PCC 7942, *Synechococcus* sp. PCC 7002, and *Synechocystis* sp. PCC 6803) were added based on their identification in Jia et al. (*82*). To exclude low quality, fragmented, or potentially non-axenic genome assemblies from further analysis, those with a CheckM completeness score <90%, a CheckM contamination score >10% or a contig N50 value <50 kb, as reported in the NCBI database were excluded. These filters left 156 high-quality Cyanobacterial genomes for further analysis.

A multispecies IsiA protein sequence (accession WP_041443458.1) was used as the query sequence in a blastp search with a minimum bit score of 250 to identify candidate IsiA proteins in the proteomes of the156 species identified above. IsiA belongs to a family of proteins that also includes Prochlorophyte chlorophyll-binding proteins (PcbA, PcbB, and PcbC), and the Chl *a*-binding proteins of Photosystem II, CP43 and CP47. CP43 proteins were excluded from the list of candidate IsiA proteins by the presence of an extended “E” loop (*83*), and CP47 genes were excluded by virtue of their lower sequence similarity (bit scores <90). From the remaining 165 candidate IsiA proteins, we could not identify any pattern that could discriminate between IsiA and PcbABC proteins. This is consistent with the conclusions of a previous study, which proposed unifying the family of IsiA and Pcb proteins and renaming them as accessory chlorophyll-binding proteins (CBPs) (*84*).

Fld proteins were identified using a multispecies FldA protein sequence (accession WP_012306911.1) in a blastp search of the 156 species’ proteomes using a minimum bit score of 100. After extensive manual curation, the candidate IsiA / Pcb / CBP proteins, the FldA / IsiB proteins, and their corresponding gene and species data were compiled into a table (Extended Dataset S3). For species with both *isiA* and *fldA* genes, the distance between genes in each species’ genome assembly was calculated with in-house scripts. Genes were considered colinear if their loci were within 5 kb of each other on the same strand of the same contig. An Euler plot demonstrating the overlap of the presence of *isiA* and *fldA* genes in each species was constructed using the eulerr package in R.

## Data Availability

The atomic coordinates for *D. salina* have been deposited in the PDB with the accession codes 9MH0 for the Fe-replete PSI-LHCI supercomplex and 9MGW for the Fe-starved PSI-LHCI_2_ supercomplex. The atomic coordinates for *D. tertiolecta* have been deposited in the PDB with the accession codes 9MH1 for the Fe-replete PSI-LHCI supercomplex and 9MGZ for the Fe-starved PSI-LHC_2_ supercomplex.

The electron microscopy maps have been deposited in the Electron Microscopy Data bank with the accession codes EMD-48266 for the Fe-replete *Dt*PSI-LHC_1_, EMD-48264 for the Fe-starved *Dt*PSI-LHC_2_ PSI, EMD-48265 for the Fe-replete *Ds*PSI-LHC_1_, and EMD-48262 for the Fe-starved *Ds*PSI-LHC_2_.

The proteomics data utilized for this dataset has been deposited in the Center for Computational Mass Spectrometry with the accession codes MSV000096479 for *D. salina* and *D. tertiolecta*, MSV000096475 for *Chla. reinhardtii*, and MSV000096362 for *Chro. zofingiensis*.

All other data are available upon request.

## Acknowledgments

The authors thank J. Oshiro for initial characterization of *Dunaliella* in Fe-depleted medium; C. Perrino for trace metal quantitation; Dr. R. Craig for bioinformatic support; D. Camacho for generating the proteomics data of *Chr. zofingiensis* in Fe-replete and Fe-starved medium; P. Tobias for computational support; Dr. D. Toso and Dr. R. Thakkar at the Cal-Cryo facility for support with cryo-EM data collection; Dr. E. Park for grids; and A. Florez for advice with data processing. This work was supported in part by the Office of Science of the US Department of Energy (DOE) grant DE-SC0020627 (to S.S.M) and through the Photosynthetic Systems program in the Office of Basic Energy Sciences (to K.K.N. and M.I.). H.W.L was supported in part by the National Science Foundation Graduate Research Fellowship Program. R.K. was supported in part by the Howard Hughes Medical Institute. Molecular graphics and analyses were performed with UCSF ChimeraX with support from NIH RO1-GM129325. (A portion of) This research was performed on a project award (<u>10.46936/lser.proj.2021.51920/60000372</u>) from the Environmental Molecular Sciences Laboratory, a DOE Office of Science User Facility sponsored by the Biological and Environmental Research program under Contract No. DE-AC05-76RL01830. E.N. and K.K.N. are investigators of the Howard Hughes Medical Institute.

## Competing interests

The author declares no competing interests.

## Author Contributions

S.S.M conceived the study.

H.W.L. and S.S.M. wrote and revised the original draft, with notable inputs and careful revisions from R.K. and M.I. All authors revised and commented on the manuscript.

H.W.L. and R.K. prepared the figures with input from all authors, especially P.G. and M.I. H.W.L., R.K., and P.G. prepared the cryo-EM samples, screened and optimized the grids.

H.W.L. and R.K. conducted the data collection, data analyses, and data processing.

H.W.L. and R.K. conducted the map interpretation, model building, and refinement with feedback from all authors.

C.D.N., S.O.P, and M.S.L. were responsible for sample preparation, data analysis and resources, respectively, for proteomics.

H.W.L. purified and biochemically analyzed the *Dunaliella* PSI-LHCI supercomplexes of *D. tertiolecta* and *D. salina* in Fe replete and Fe starvation medium.

M.I., R.K., P.G., and S.S.M trained and supervised H.W.L.

S.D.G. identified *isiA* and *isiB* orthologs in cyanobacteria.

S.S.M., K.K.N., and E.N. provided resources, infrastructure and feedback.

## Extended Data

**Extended Data Fig. 1.**
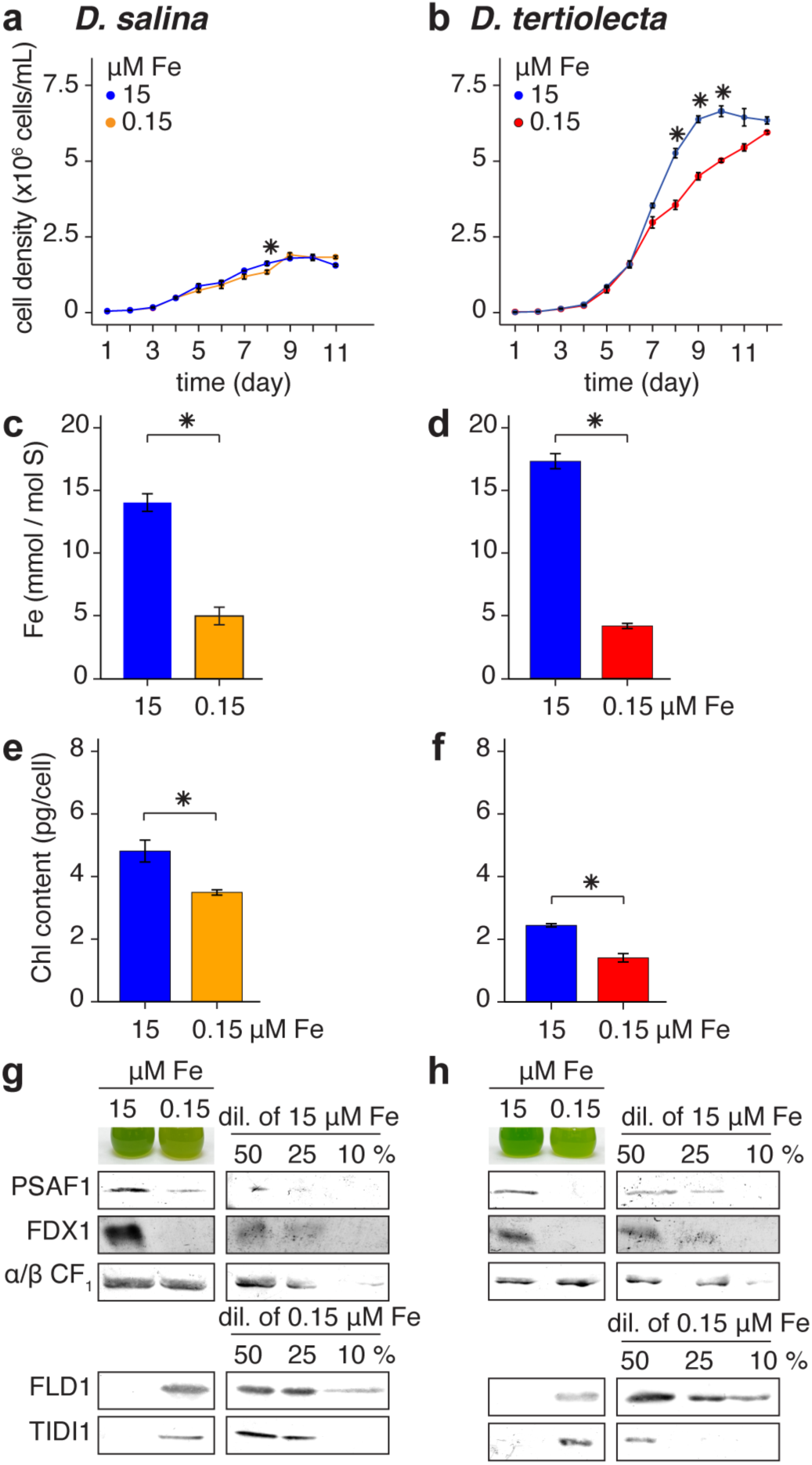
*Dunaliella* spp. are resilient to Fe starvation. **a**,**b**, Growth, **c**,**d**, Fe content, **e**,**f**, Chl content, and **g**,**h**, immunodetection of photosynthetic proteins in *Dunaliella salina* (left) and *Dunaliella tertiolecta* (right) grown in Fe-replete (blue) and Fe-depleted (orange for *D. salina*, red for *D. tertiolecta*) medium. Standard error based on three independent cultures. *Asterisk* (*) denotes statistically significant difference relative to 15 µM Fe (*p* value < 0.05).

**Extended Data Fig. 2.**
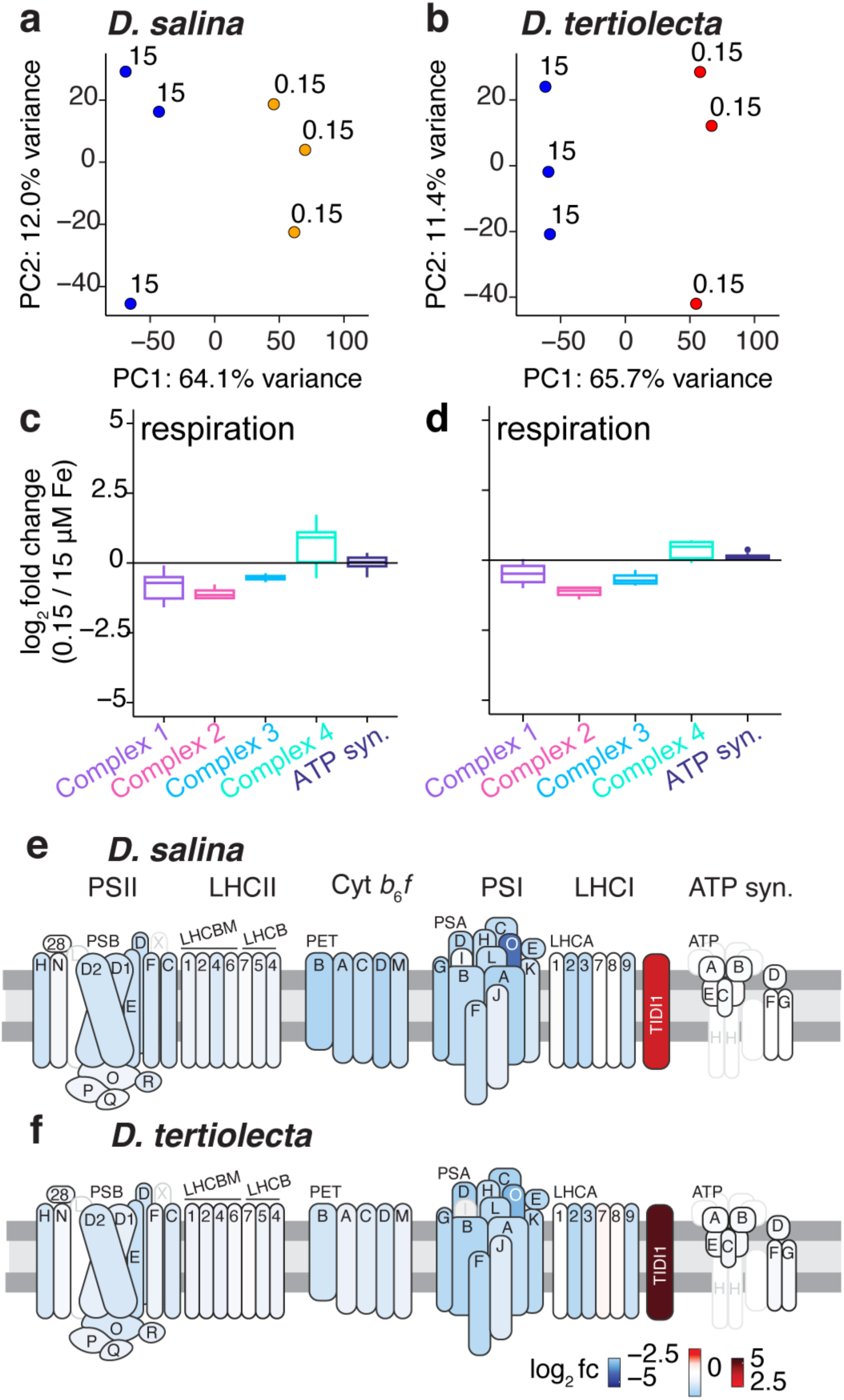
The *Dunaliella* proteome responds to Fe-depletion. **a**,**b**, The principal component analysis (PCA) separates the proteome of both *D. salina* (**a**) and *D. tertiolecta* (**b**) based on Fe availability (15 µM versus 0.15 µM medium). **c**,**d**, The log_2_ fold change of components of respiration (Complex 1, Complex 2, Complex 3, Complex 4, and ATP synthase). The full list of proteins included in the analysis is found in Extended Data 2. **e**,**f**, Overview of changes in abundance of proteins of the photosynthetic electron transfer change. Relative changes in protein abundances in Fe-starved (0.15 µM Fe) versus Fe-replete (15 µM Fe) medium as log_2_ fold changes (red, increase; blue, decrease). Proteins are schematically assembled according to their position within the respective complex.

**Extended Data Fig. 3.**
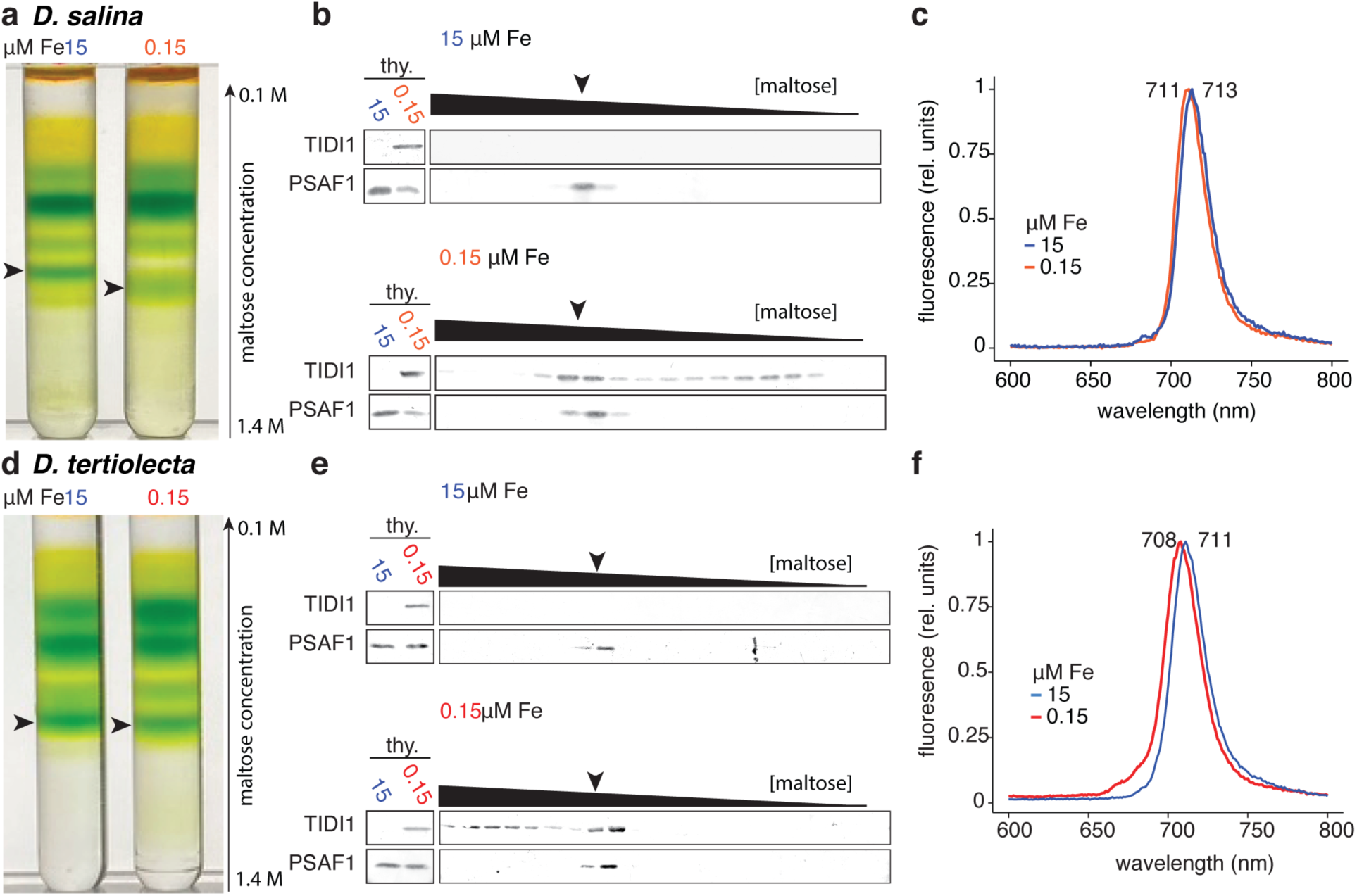
Biochemical characterization of *Dunaliella* spp. PSI-LHCI supercomplex in Fe -replete and -starved conditions. **a**,**d**, Isolation of PSI from thylakoid membranes of cells grown either in Fe-replete (left, 15 µM Fe) or -starved (right, 0.15 µM Fe) medium using maltose density gradient centrifugation for *D. salina* (**a**) and *D. tertiolecta* (**d**). Arrows indicate the locations of the bands corresponding to PSI. A representative result was shown from three independent thylakoid membrane isolations. **b**,**e**, Immunoblot analysis showing the presence of PSI and TIDI1 in the thylakoids (thy.) (left) and location of PSI and TIDI1 within the maltose density gradient (right) from cells grown in either 15 µM Fe (top) or 0.15 µM Fe (bottom) medium for *D. salina* (**b**) and *D. tertiolecta* (**e**). **c**,**f**, 77 K chlorophyll fluorescence emission spectra of isolated PSI isolated from cells grown 15 µM Fe (blue) or 0.15 µM Fe (orange) medium for *D. salina* (**c**) and *D. tertiolecta* (**f**).

**Extended Data Fig. 4.**
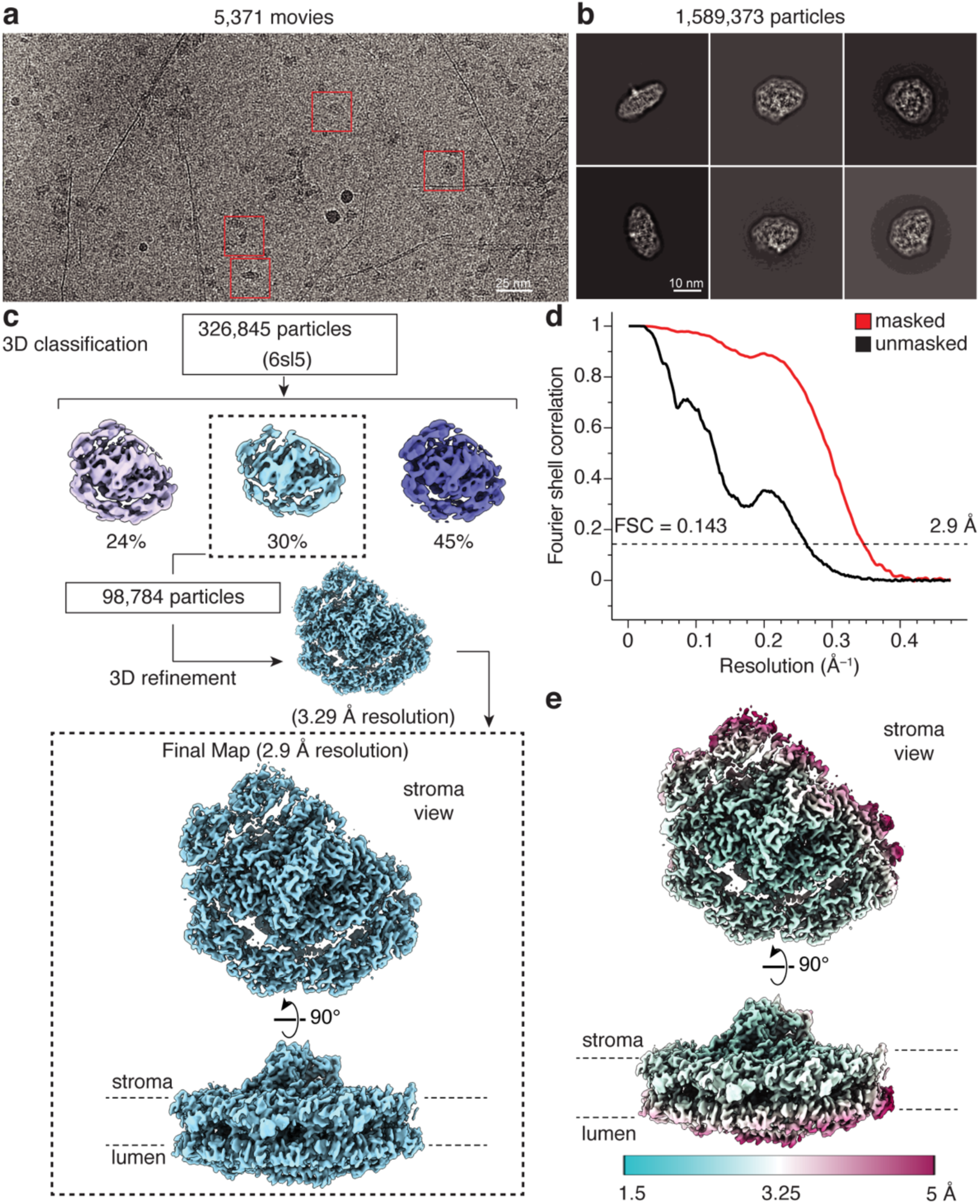
Processing flowchart for the *D. salina* PSI-LHCI supercomplex in Fe- replete conditions. **a**, Representative cryo-EM image of *D. salina* Fe-replete PSI-LHCI supercomplexes. **b**, Representative 2D class averages of *D. salina* Fe-replete PSI-LHCI supercomplexes. Box size is 240 Å. **c**, Processing flowchart for the cryo-EM dataset using RELION (v.3.0). **d**, FSC curve of the final reconstruction. The profiles for masked and unmasked are red and black, respectively. **e**, Local resolution for the final cryo-EM map.

**Extended Data Fig. 5.**
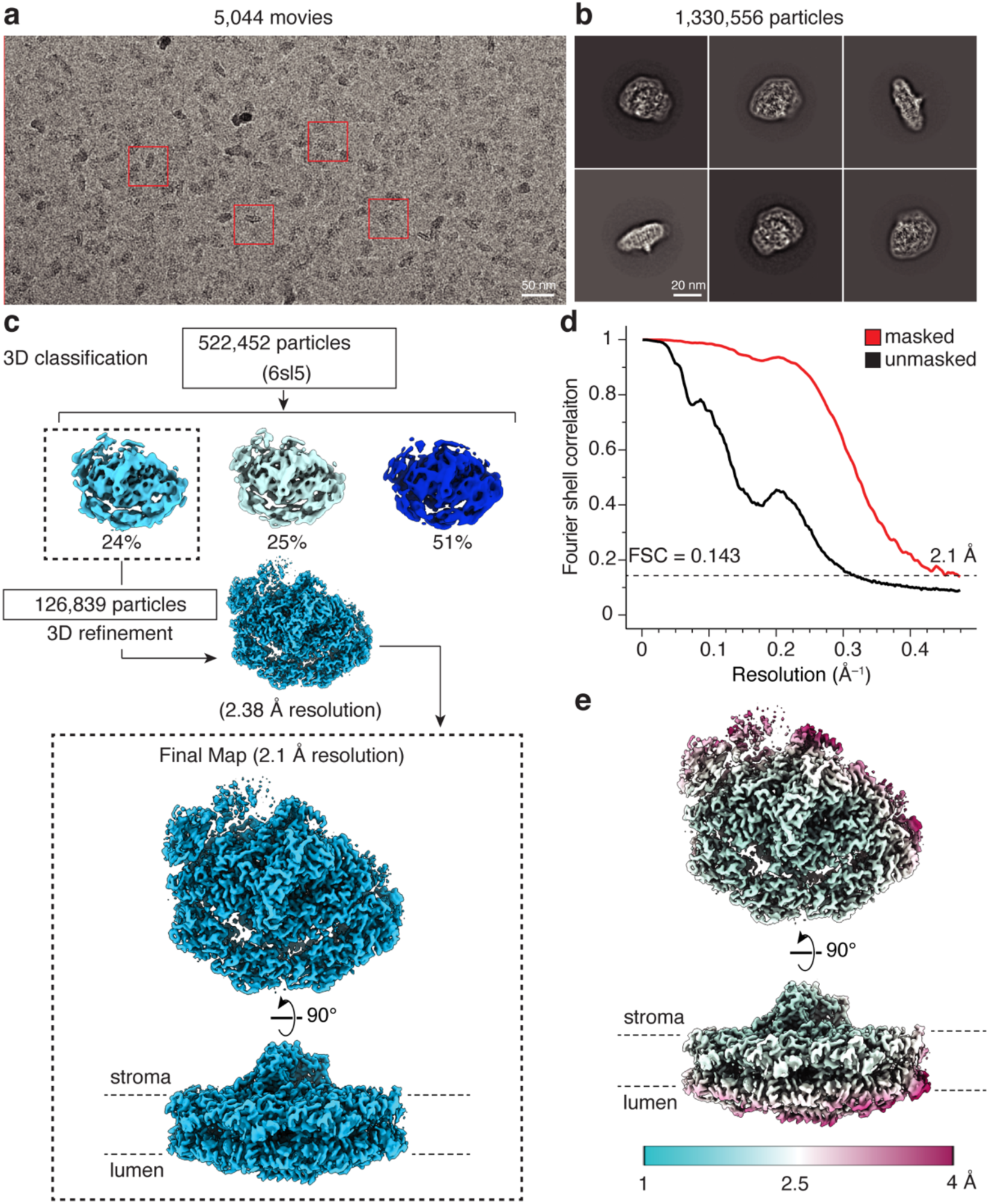
Processing flowchart for the *D. tertiolecta* PSI-LHCI supercomplex in Fe-replete conditions. **a**, Representative cryo-EM image of *D. tertiolecta* Fe-replete PSI-LHCI supercomplexes. **b**, Representative 2D class averages of *D. tertiolecta* Fe-replete PSI-LHCI supercomplexes. Box size is 240 Å. **c**, Processing flowchart for the cryo-EM dataset with RELION (v.3.0). **d**, FSC curve of the final reconstruction. The profiles for masked and unmasked are red and black, respectively. **e**, Local resolution for the final cryo-EM map.

**Extended Data Fig. 6.**
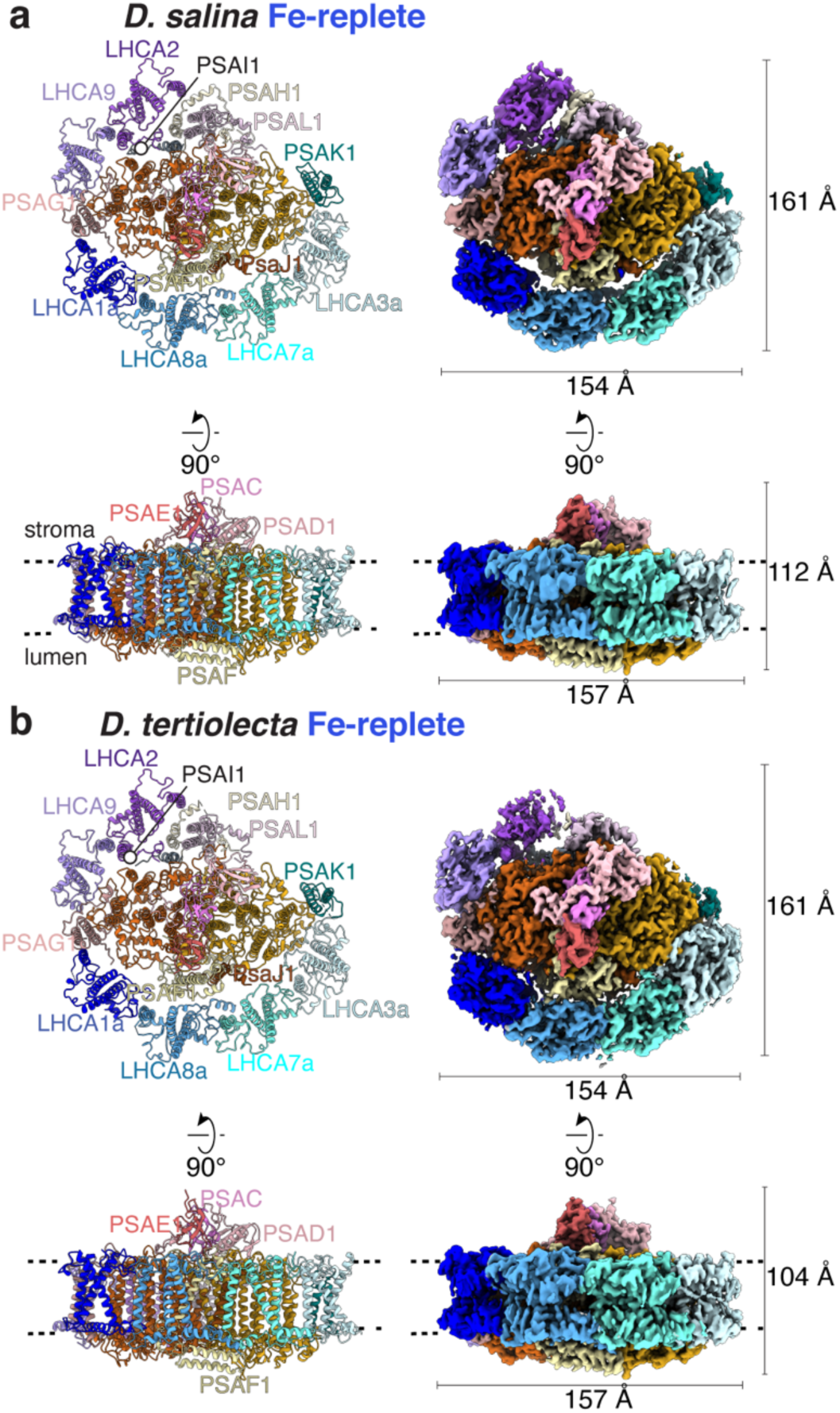
Structural overview of the *Dunaliella* spp. PSI-LHCI_1_ supercomplexes from Fe-replete cells. **a**,**b**, The model (left) and the cryo-EM density maps (right) for PSI-LHCI_1_ supercomplexes from Fe-replete *Dunaliella salina* (**a**) and *Dunaliella tertiolecta* (**b**).

**Extended Data Fig. 7.**
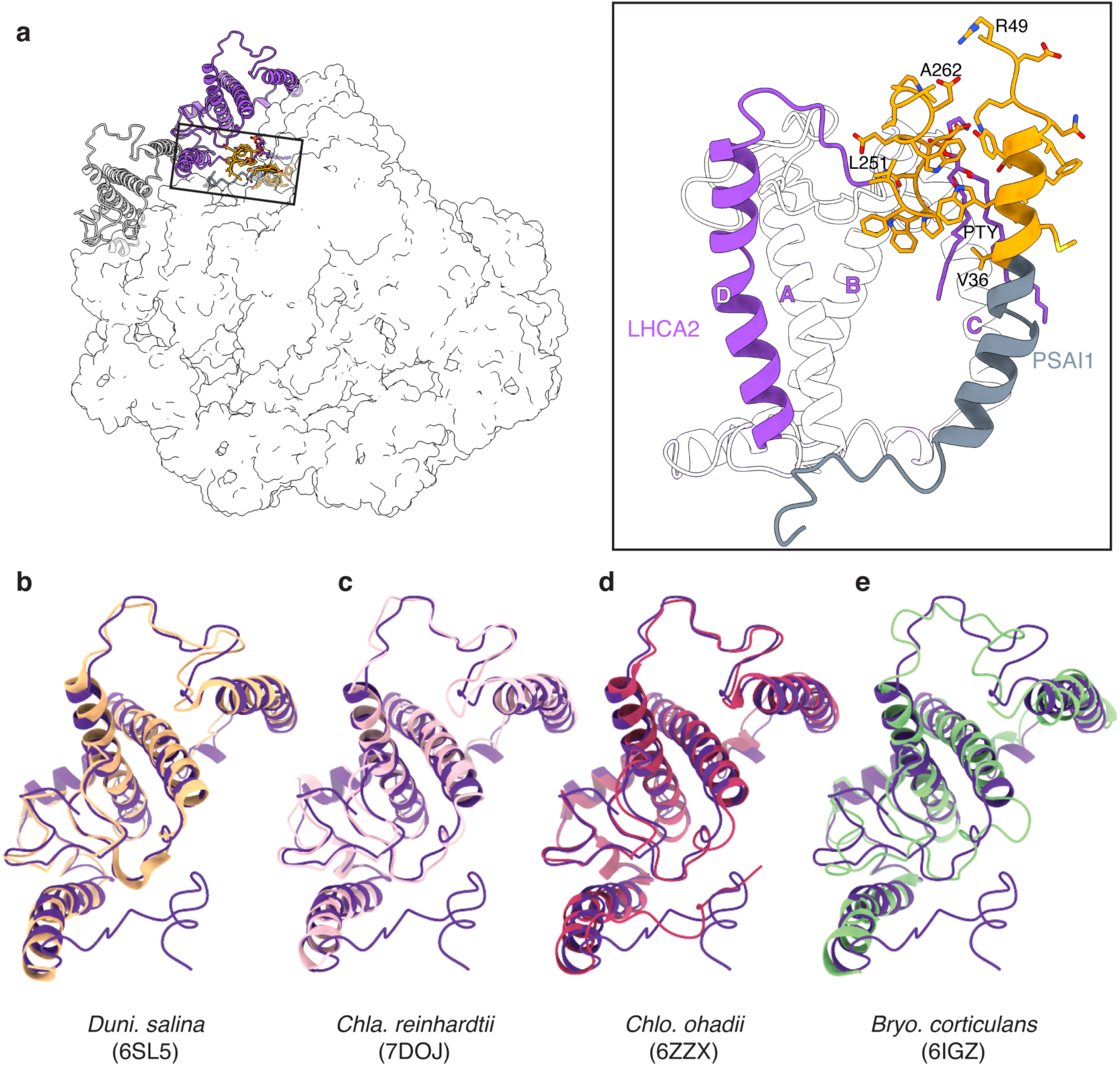
*Ds*PSI-LHCI_1_ LHCA2 has an extended C-terminus. **a**, Left: Model of *D. salina* LHCA2 (purple) with the extended C-terminus with PSAI1 (gray) and LHCA9 (white). Right: The extended C-terminus of LHCA2 interacts with PSAI1. Residues involved in the interaction are shown in orange stick representations. **b**–**e**, Overlay of *Ds*LHCA2 (purple) and previously reported models (**b**) *Duni. salina* (yellow, 6SL5), (**c**) *Chla. reinhardtii* (pink, 7D0J), (**d**) *Chlo. ohadii* (red, 6ZZX), and (**e**) *Bryo. corticulans* (green, 6IGZ) LHCA2.

**Extended Data Fig. 8.**
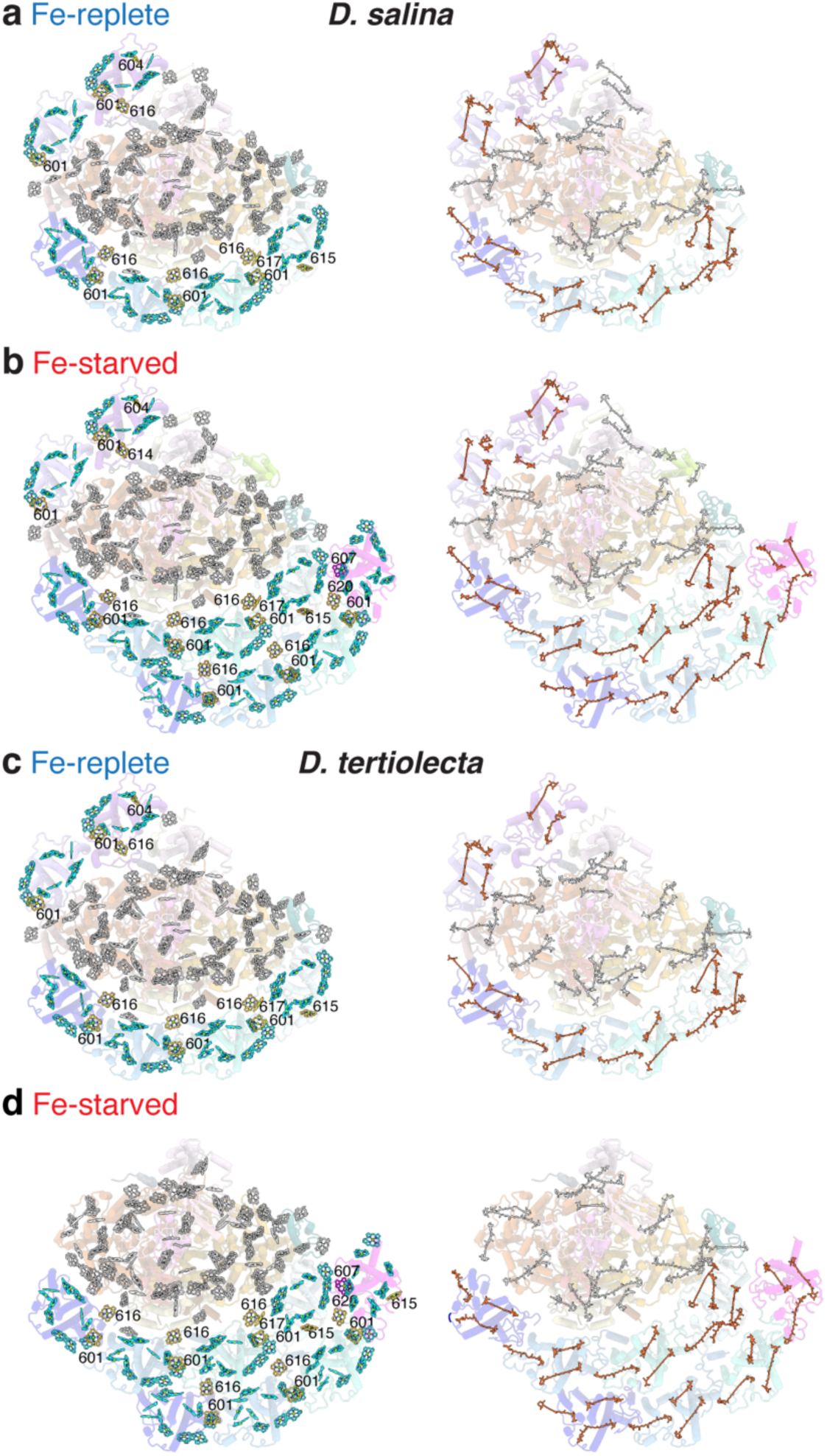
Pigment arrangement in the *Ds*PSI-LHCI and *Dt*PSI-LHCI supercomplexes. **a-d**, Left: Chl arrangement in *Ds*PSI-LHCI_1_ (**a**), *Ds*PSI-LHCI_2_ (**b**), *Dt*PSI-LHCI_1_ (c) and *Dt*PSI-LHCI_2_ (**d**) supercomplexes. Chls in the PSI core are colored in gray. Conserved Chl sites 601-614 are colored in cyan, and specific Chl sites 613-620 are colored in orange. **a-d**, Right: Car arrangement in *Ds*PSI-LHCI_1_ (**a**), *Ds*PSI-LHCI_2_ (**b**), *Dt*PSI-LHCI_1_ (**c**) and *Dt*PSI-LHCI_2_ (d) supercomplexes. The PSI core Cars are colored in gray, and Car found in LHCA subunits are colored in red.

**Extended Data Fig. 9.**
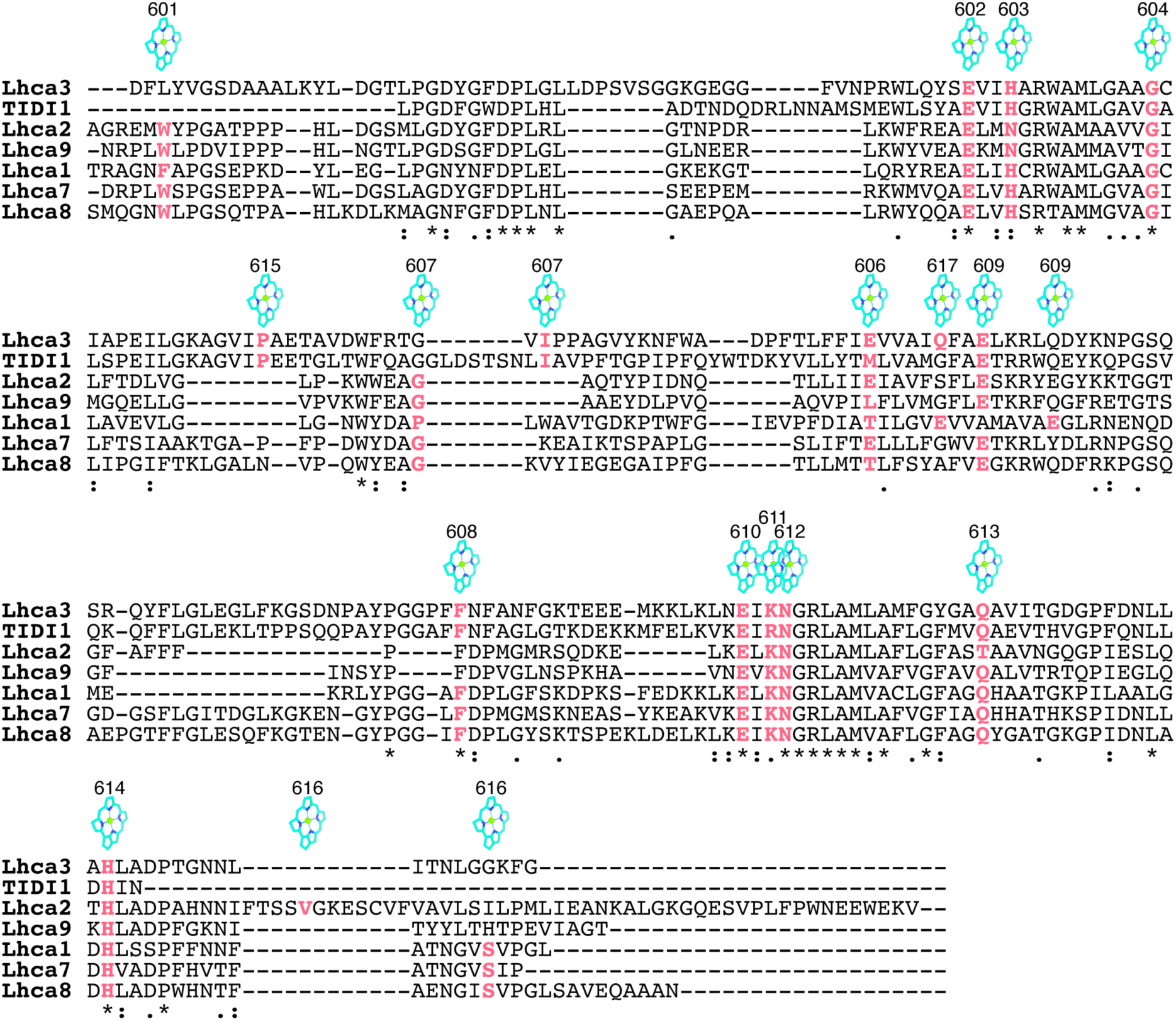
Chl coordination of LHCI subunits from *D. salina* PSI-LHCI supercomplex. Multiple sequence alignment of LHCI subunits with residues involved in Chl coordination or contributing to Chl coordination are shaded in blue. Chl and their annotations are based on PDB:1RWT and shown above.

**Extended Data Fig. 10.**
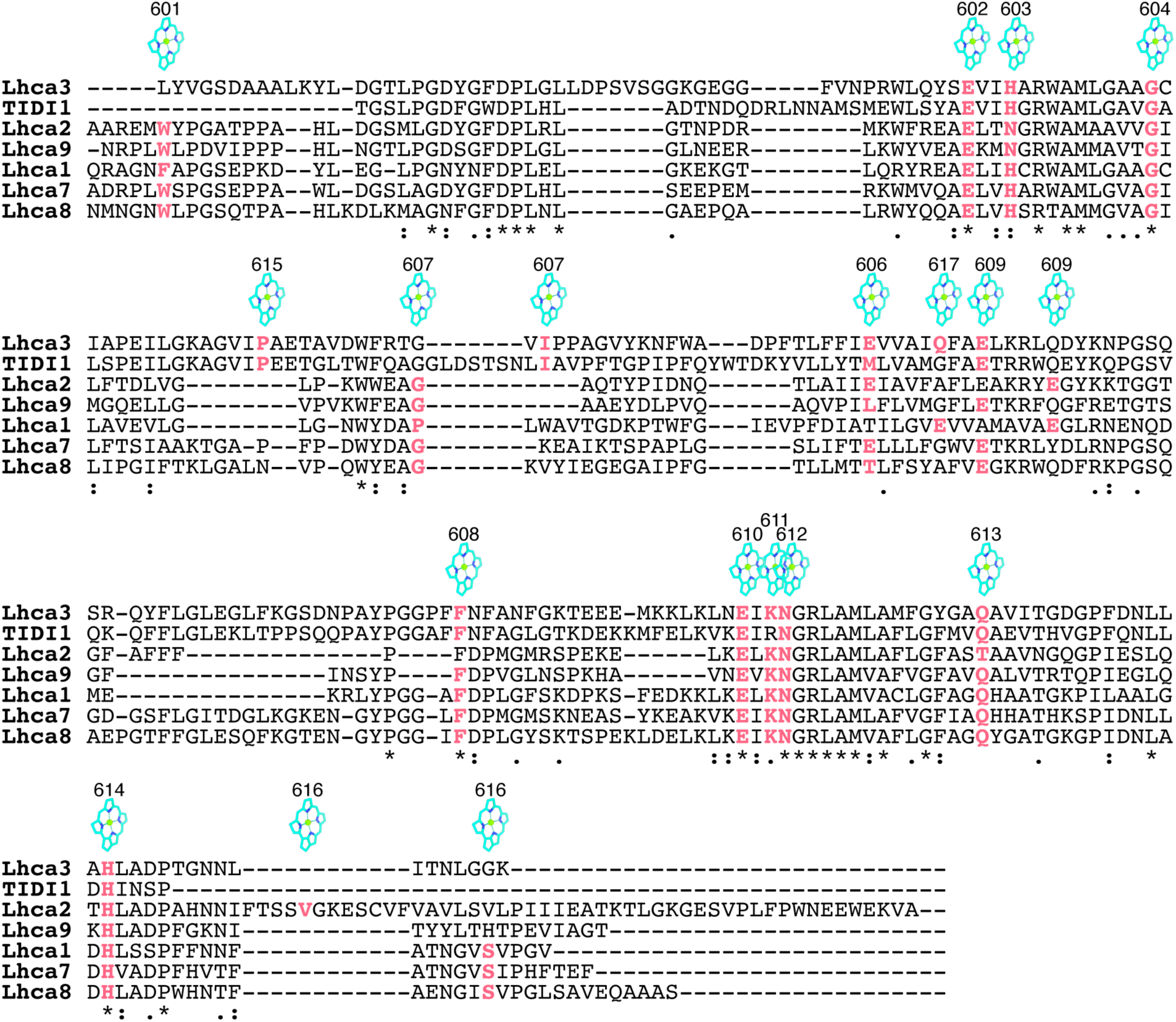
Chl coordination of LHCI subunits from *D. tertiolecta* PSI-LHCI supercomplex. Multiple sequence alignment of LHCI subunits with residues involved in Chl coordination or contributing to Chl coordination are shaded in blue. Chl and their annotations are based on PDB:1RWT and shown above.

**Extended Data Fig. 11.**
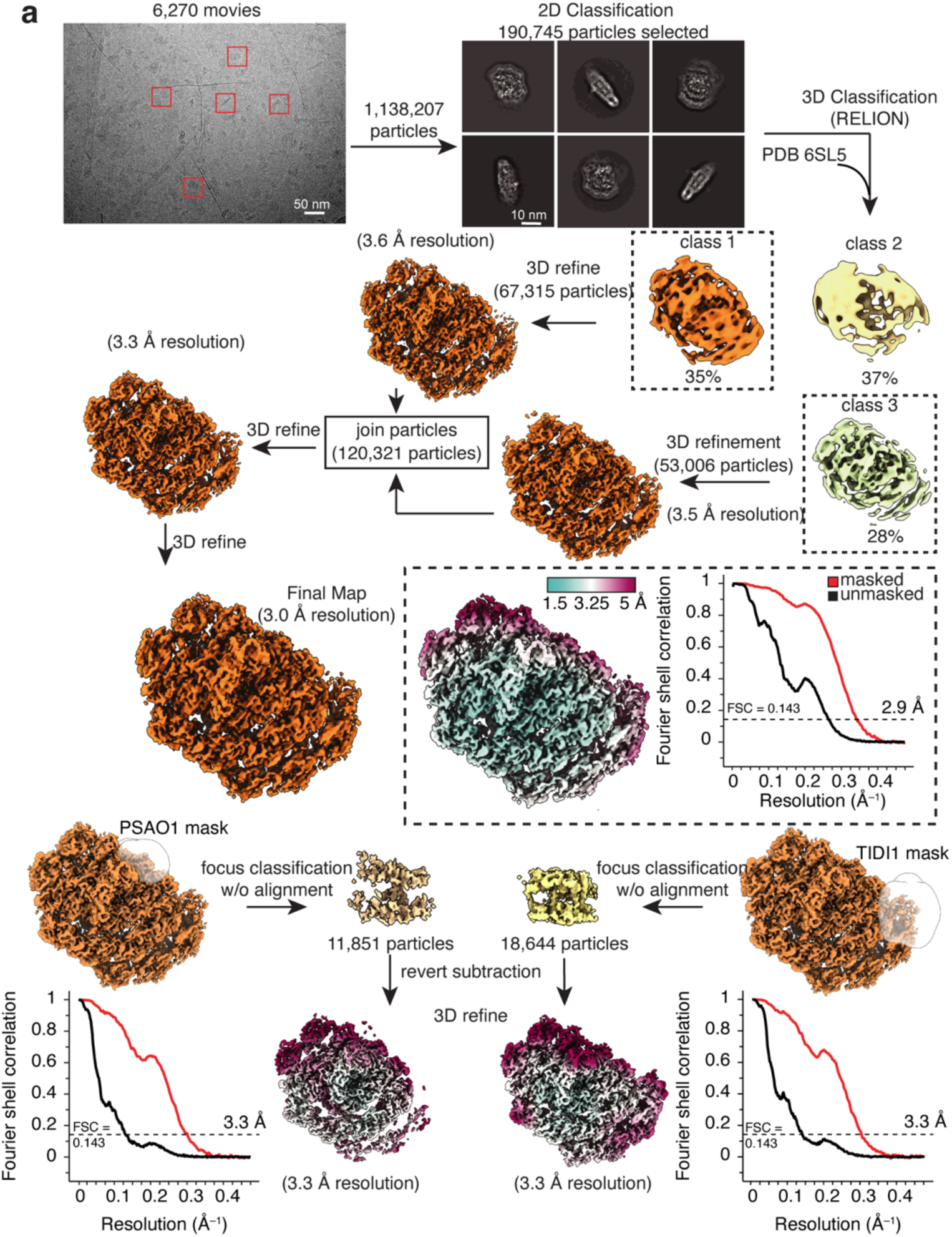
Processing flowchart for *D. salina* PSI-LHCI supercomplex in Fe-starved conditions. **a**, Flowchart showing all 3D classification and refinement steps used to determine of the overall Fe-starved *D. salina* PSI using RELION (v.3.0).

**Extended Data Fig. 12.**
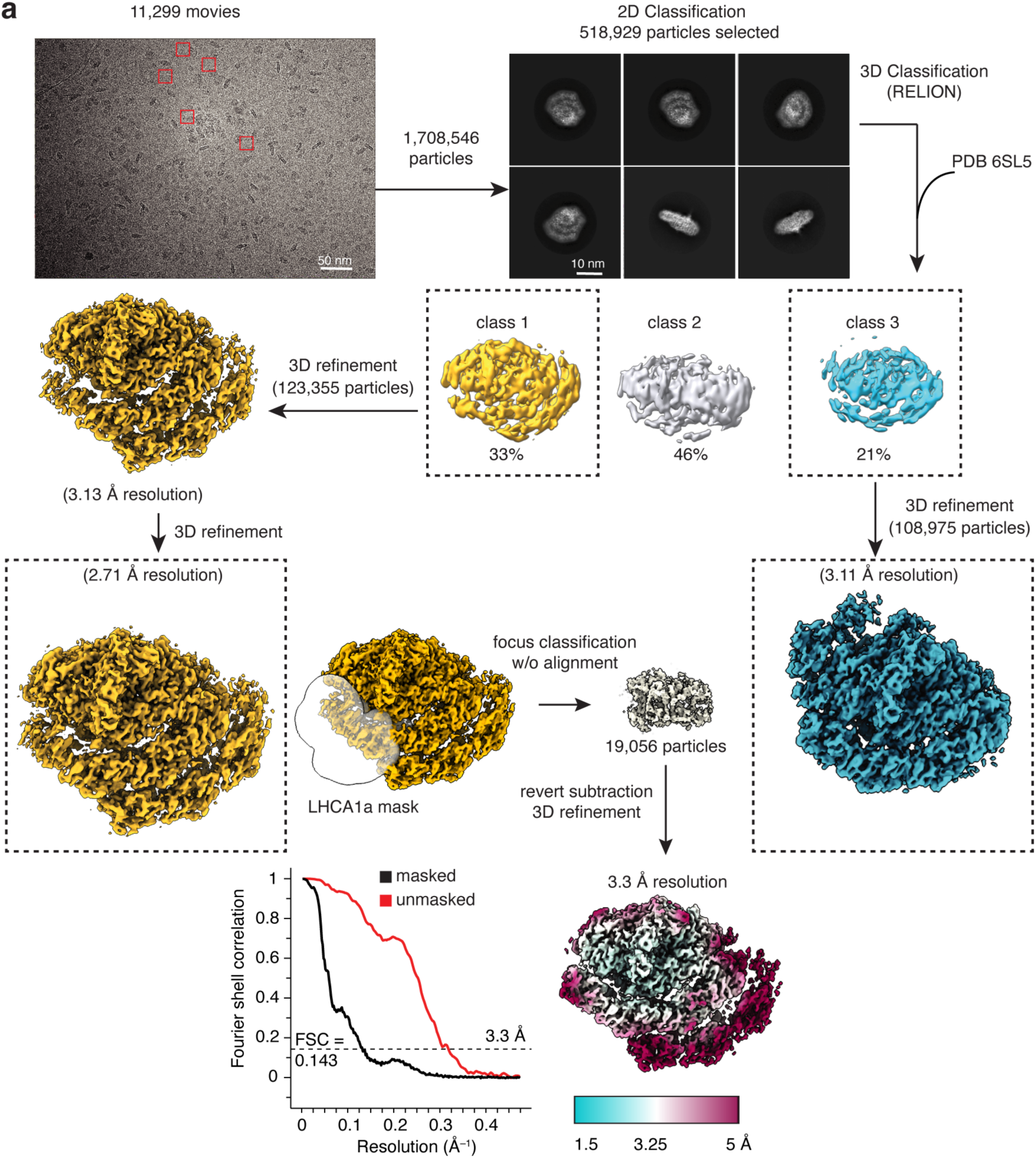
Processing flowchart for *D. tertiolecta* PSI-LHCI supercomplex in Fe-starved conditions. **a**, Flowchart showing all 3D classification and refinement steps used to determine of the overall Fe-starved *D. tertiolecta* PSI using RELION (v.3.0).

**Extended Data Fig. 13.**
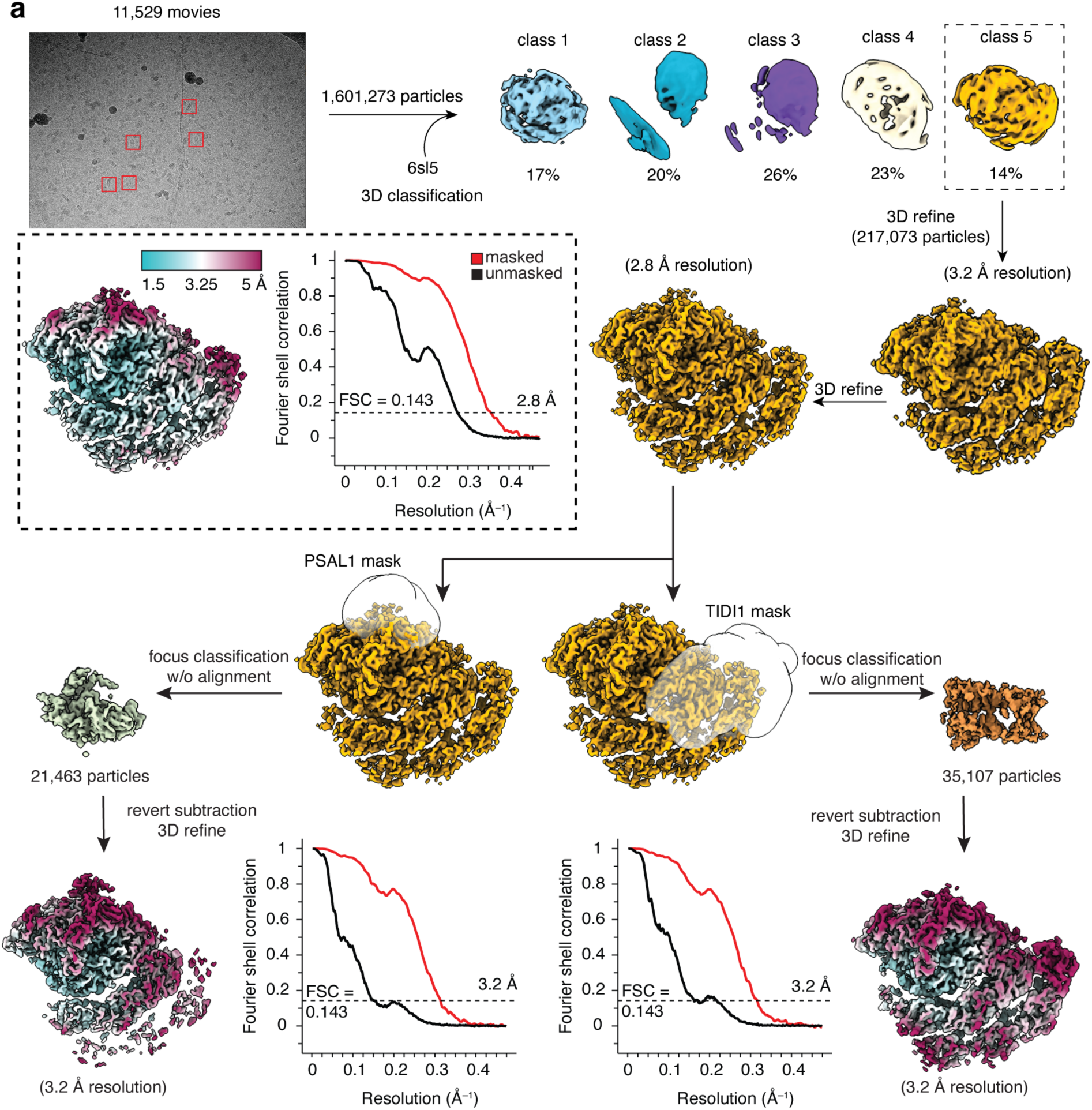
Processing flowchart for *D. tertiolecta* PSI-LHCI supercomplex in Fe-starved conditions isolated by the GraFix strategy. **a**, Flowchart showing all 3D classification and refinement steps used to determine of the overall Fe-starved *D. tertiolecta* PSI using RELION (v.3.0).

**Extended Data Fig. 14.**
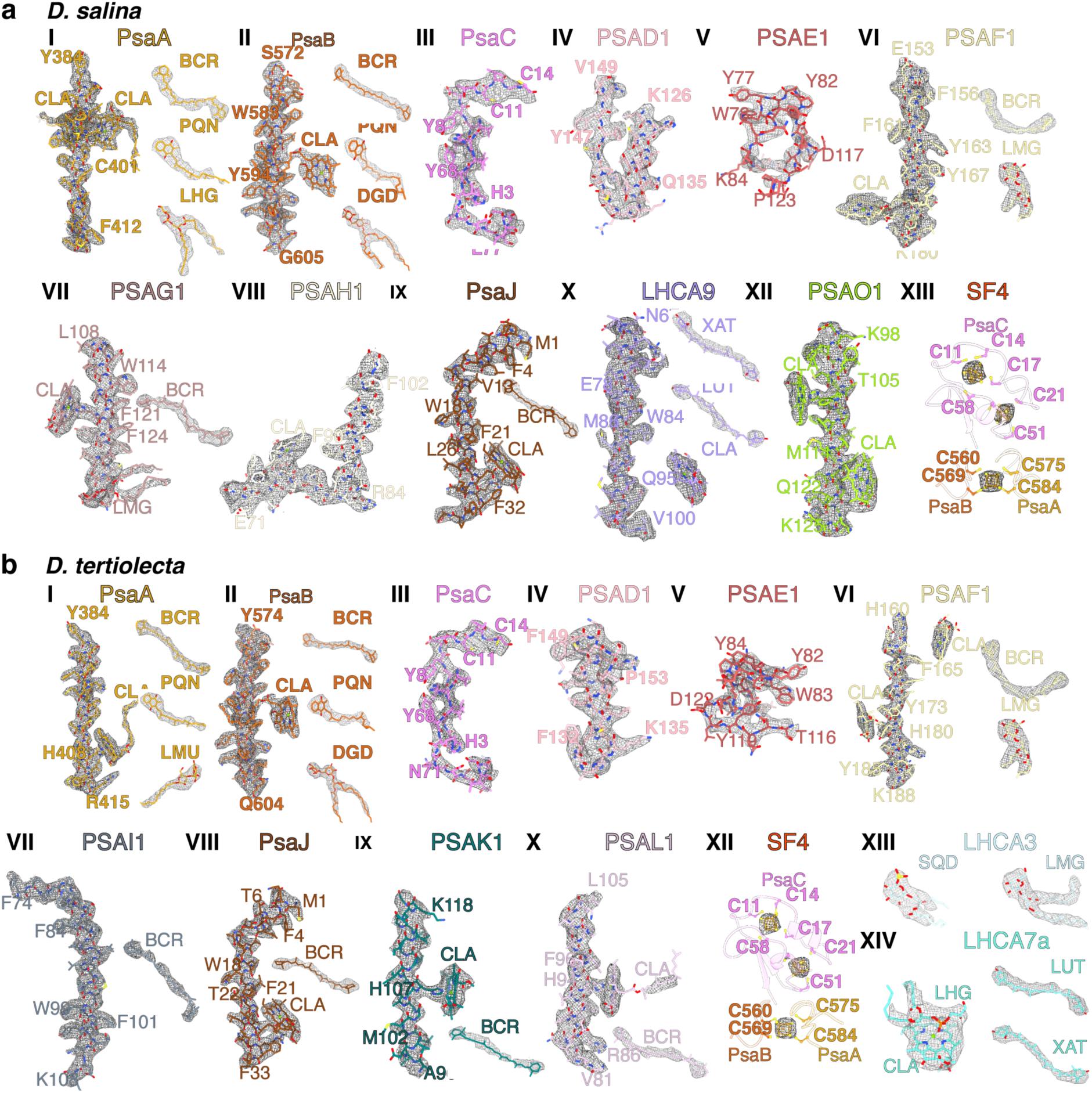
Model fitting of the *Dunaliella* spp. PSI-LHCI_2_ supercomplexes. **a**,**b**, Model fitting of the PSI-LHCI_2_ supercomplexes in the cryo-EM density map for different subunits of the structure in *D. salina* (**a**) and *D. tertiolecta* (**b**).

**Extended Data Fig. 15.**
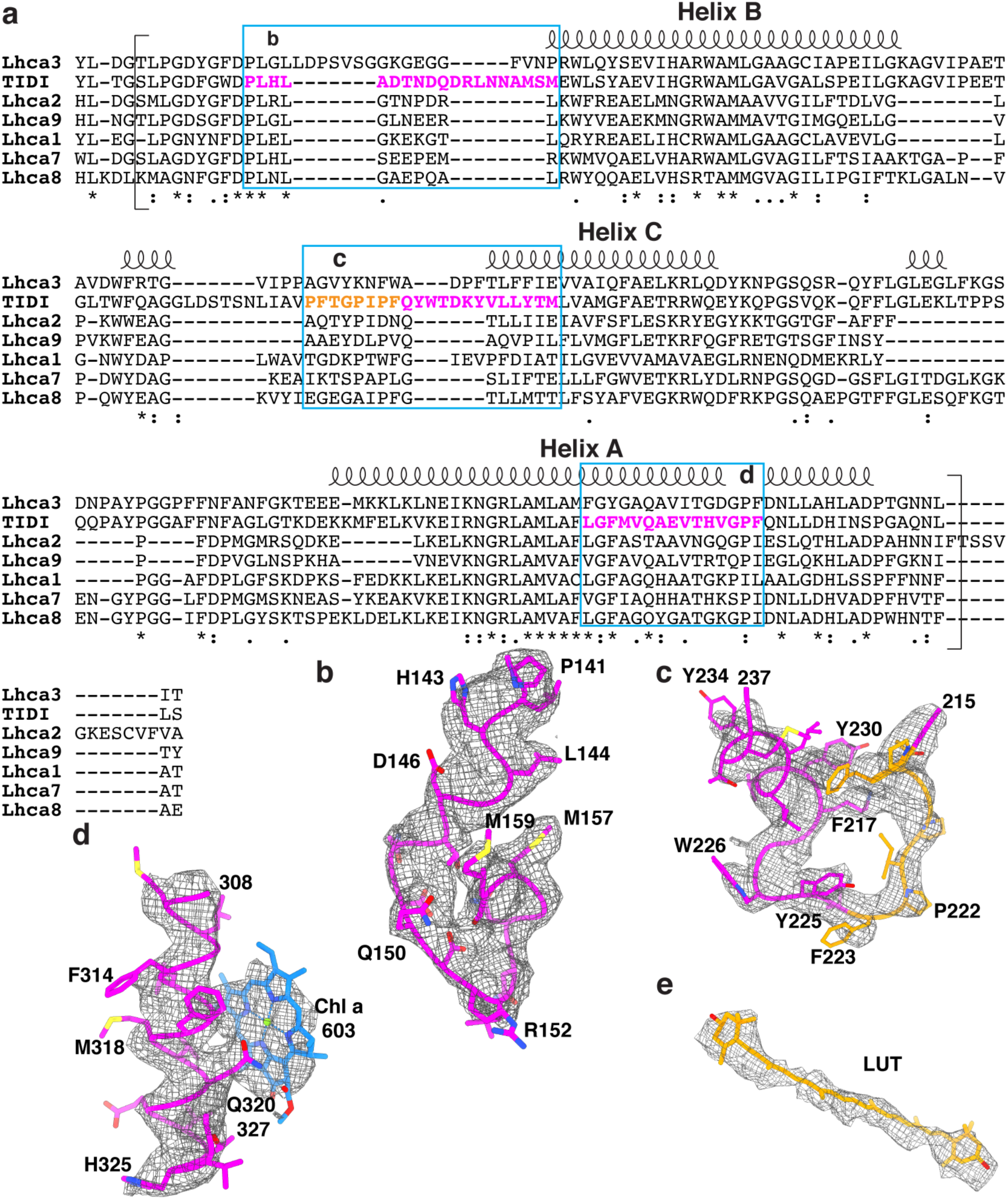
Molecular details of *D. salina* TIDI1. **a**, Multiple sequence alignment of *D. salina* LHCI subunits and TIDI1. The secondary structure is shown above the sequences. TIDI1 sequences best matching the cryo-EM density maps are shaded in magenta and boxed in blue frames. The amino acids in the conserved TIDI1 motif (PFXGX_2_PF) are shaded in orange. **b**-**f**, Cryo-EM density maps and structures of TIDI1 in regions best matching the TIDI1 sequences and pigment molecules.

**Extended Data Fig. 16.**
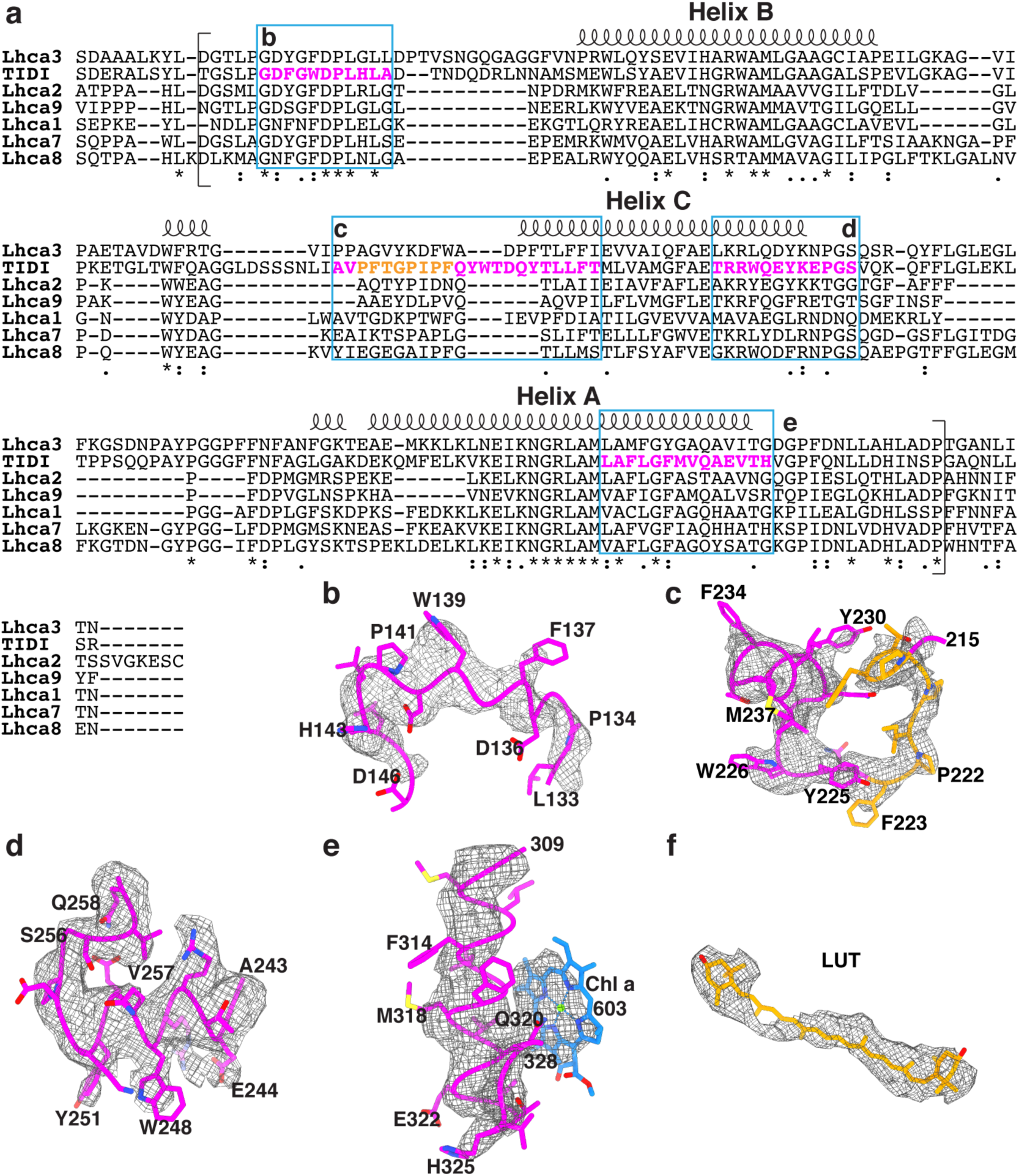
Molecular details of *D. tertiolecta* TIDI1. **a**, Multiple sequence alignment of *D. tertiolecta* LHCI subunits and TIDI1. The secondary structure is shown above the sequences. TIDI1 sequences best matching the cryo-EM density maps are shaded in magenta and boxed in blue frames. The conserved TIDI1 motif are shaded in orange. **b**-**f**, Cryo-EM density maps and structures of TIDI in regions best matching the TIDI1 sequences and pigment molecules.

**Extended Data Fig. 17.**
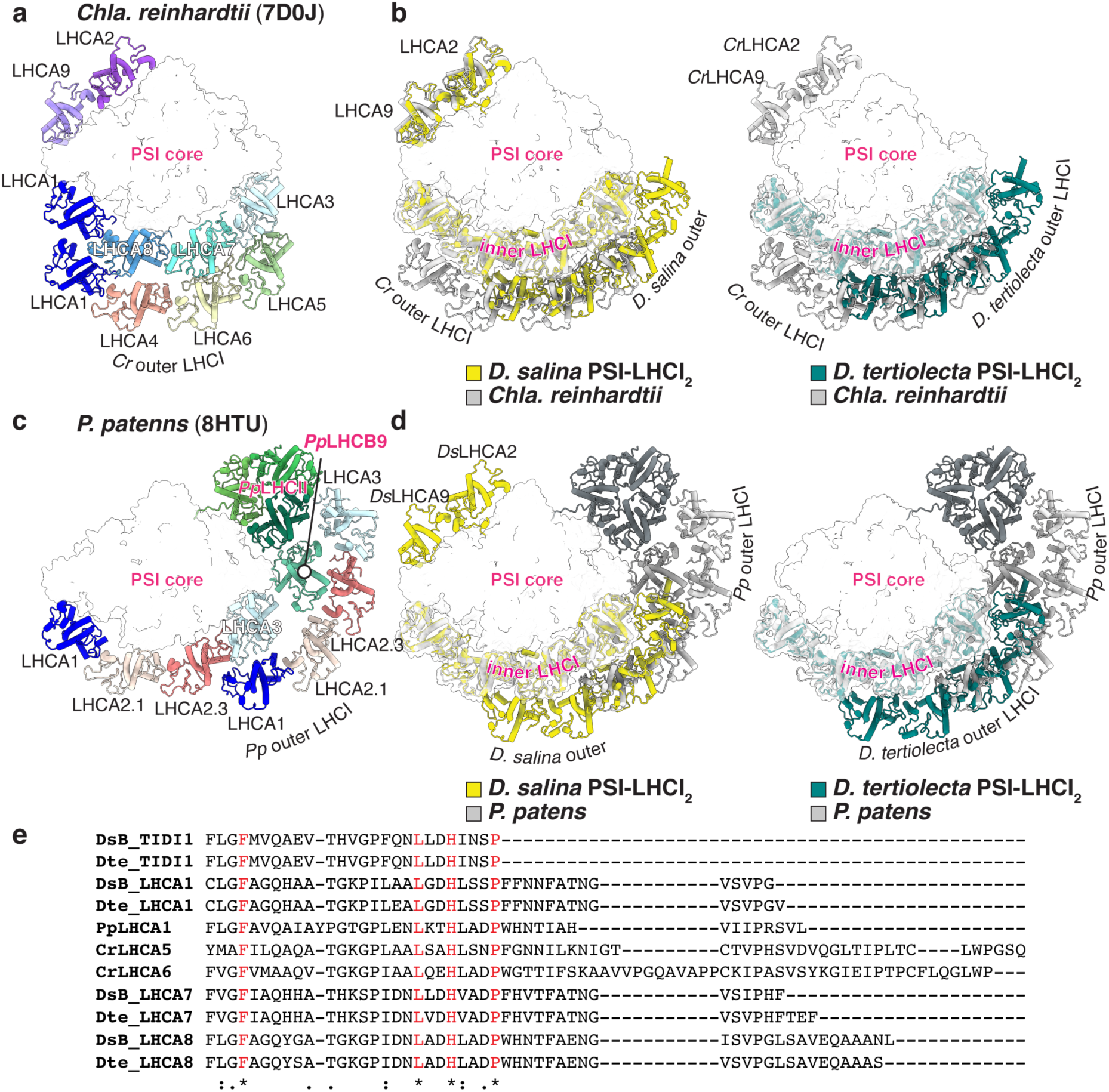
Comparison of LHCI tetramers from *Dunaliella* spp. compared to Chla. reinhardtii *and* Phys. Patens. **a**,**c**, Stromal view of *Cr*PSI-LHCI (PDB code 7D0J) (**a**) and large *Pp*PSI-LHCI (PDB code 8HTU) (**c**) supercomplex. **b**,**d**, *Ds*PSI-LHCI_2_ supercomplex (yellow, left) and *Dt*PSI-LHCI_2_ supercomplex (teal, right) superimposed on *Cr*PSI-LHCI (gray) (**b)** or on *Pp*PSI-LHCI (gray) (**d**). The PSI core is shown either as cylinders (left) or surface view (middle, right). LHC subunits are shown as cylinders. **e**, Multiple sequence alignment results of the C-terminal ends of the outer tetramer LHCs for *Dunaliella* spp., *P. patens* LHCA1, and *Chla. reinhardtii* LHCA5 and LHCA6.

**Extended Data Fig. 18.**
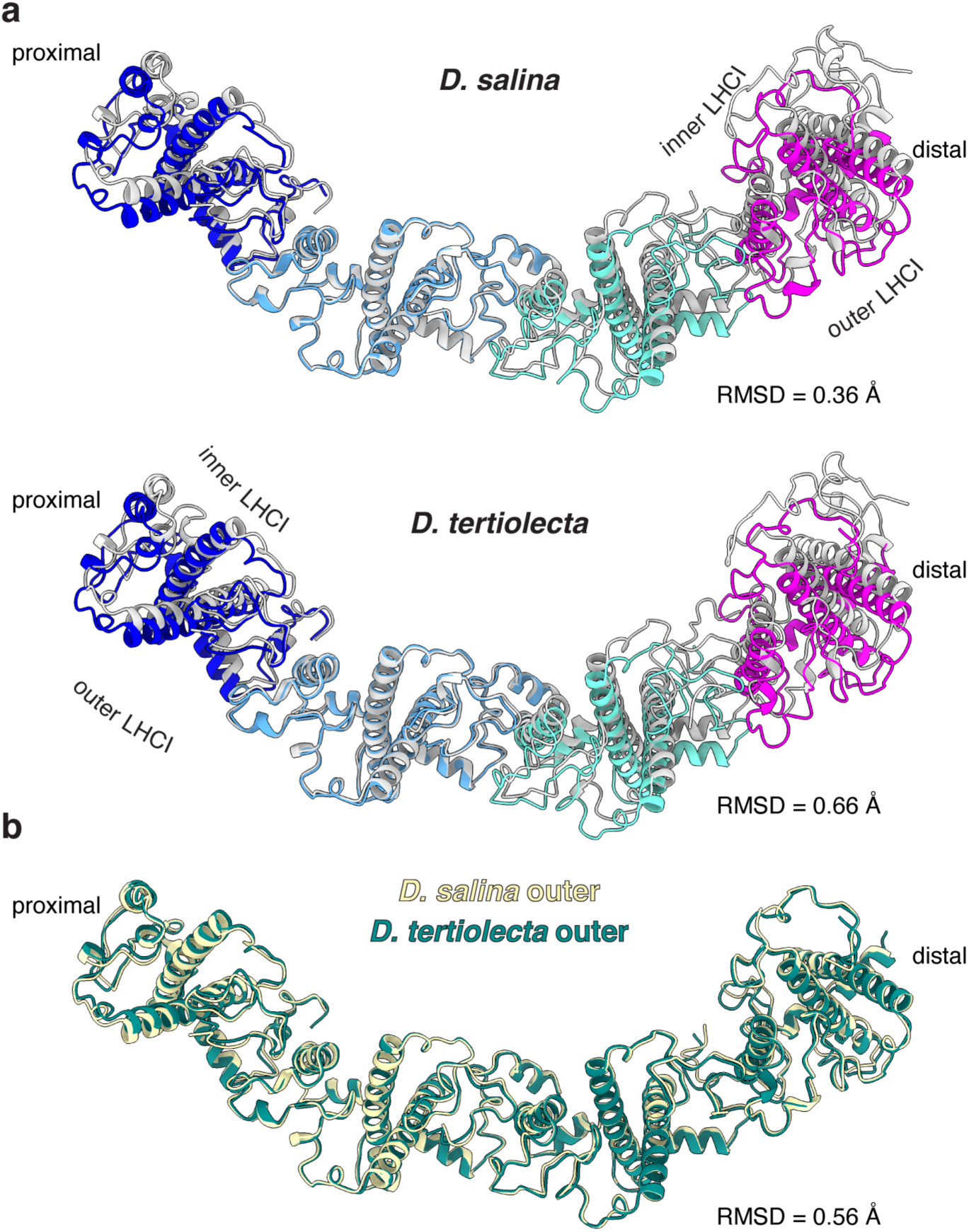
Comparison of LHCI tetramers from *Dunaliella* spp. **a**, Superimposition of inner LHCI tetramer (gray and LHCA3 as light blue) and outer LHCI tetramer (blue colored and TIDI1 as magenta) of the *Ds*PSI-LHCI_2_ supercomplex (top) and *Dt*PSI-LHCI_2_ supercomplex (bottom). **b**, Superimposition of the outer LHCI tetramer of *Ds*PSI-LHCI_2_ supercomplex (yellow) and *Dt*PSI-LHCI_2_ supercomplex (teal).

**Extended Data Fig. 19.**
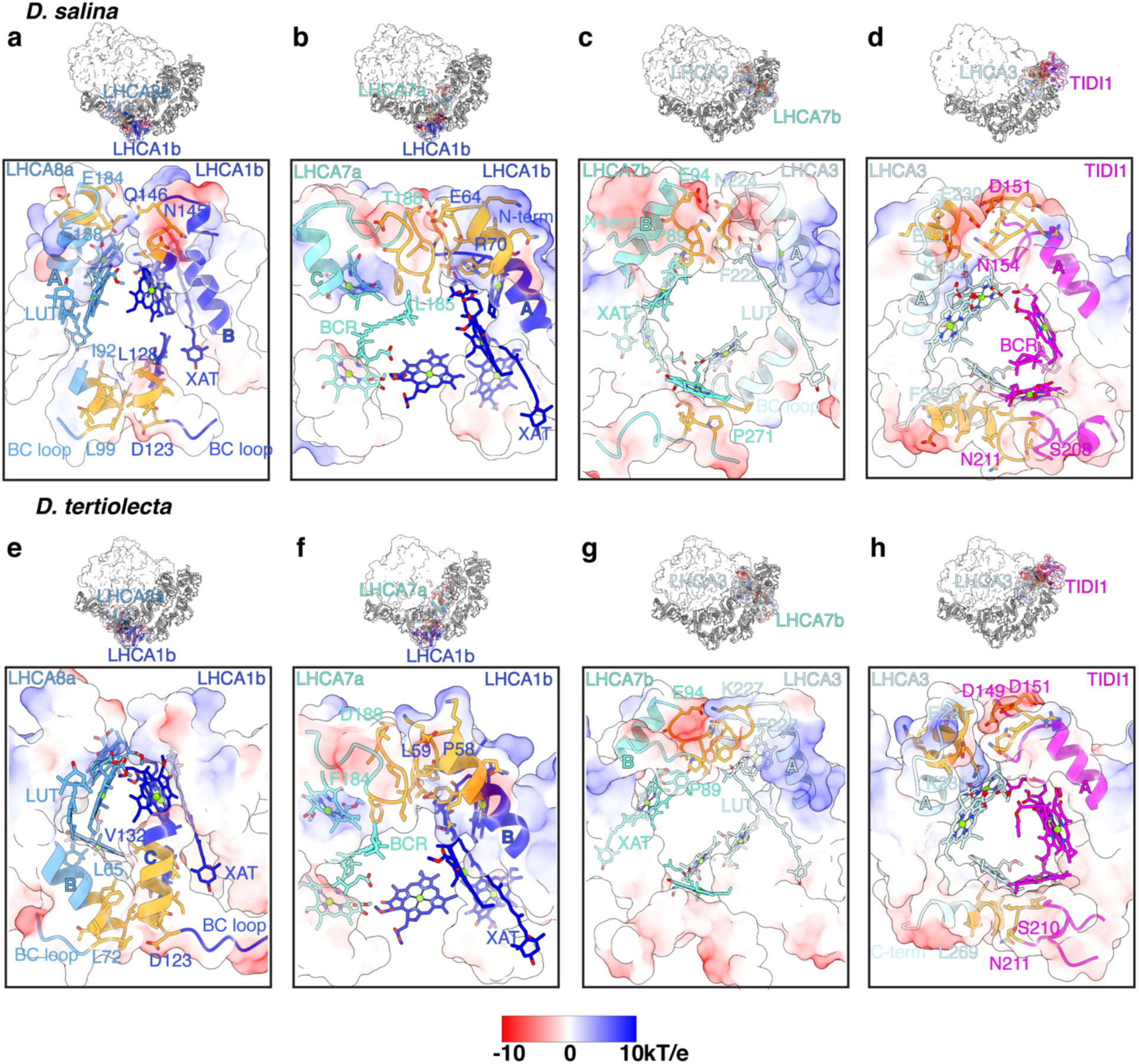
Interactions between the LHC tetramers of the *Dunaliella* spp. PSI- LHCI_2_ supercomplexes. Electrostatic potential surface representation of the proposed interaction sites between the inner and outer LHC tetramers of the (**a**-**d**) *Dunaliella salina* and (**e**-**h**) *Dunaliella tertiolecta* PSI-LHCI_2_ supercomplexes from Fe-starved conditions. Positively charged side chains are shown in blue and negatively charged residues are shown in red as indicated by the scale bar (-10 to 10 kT/e). The columbic potentials were computed using AMBER built-in Chimera X-1.7.1. Residues involved in the interaction between the outer and inner LHCI tetramer are highlighted in orange.

**Extended Data Fig. 20.**
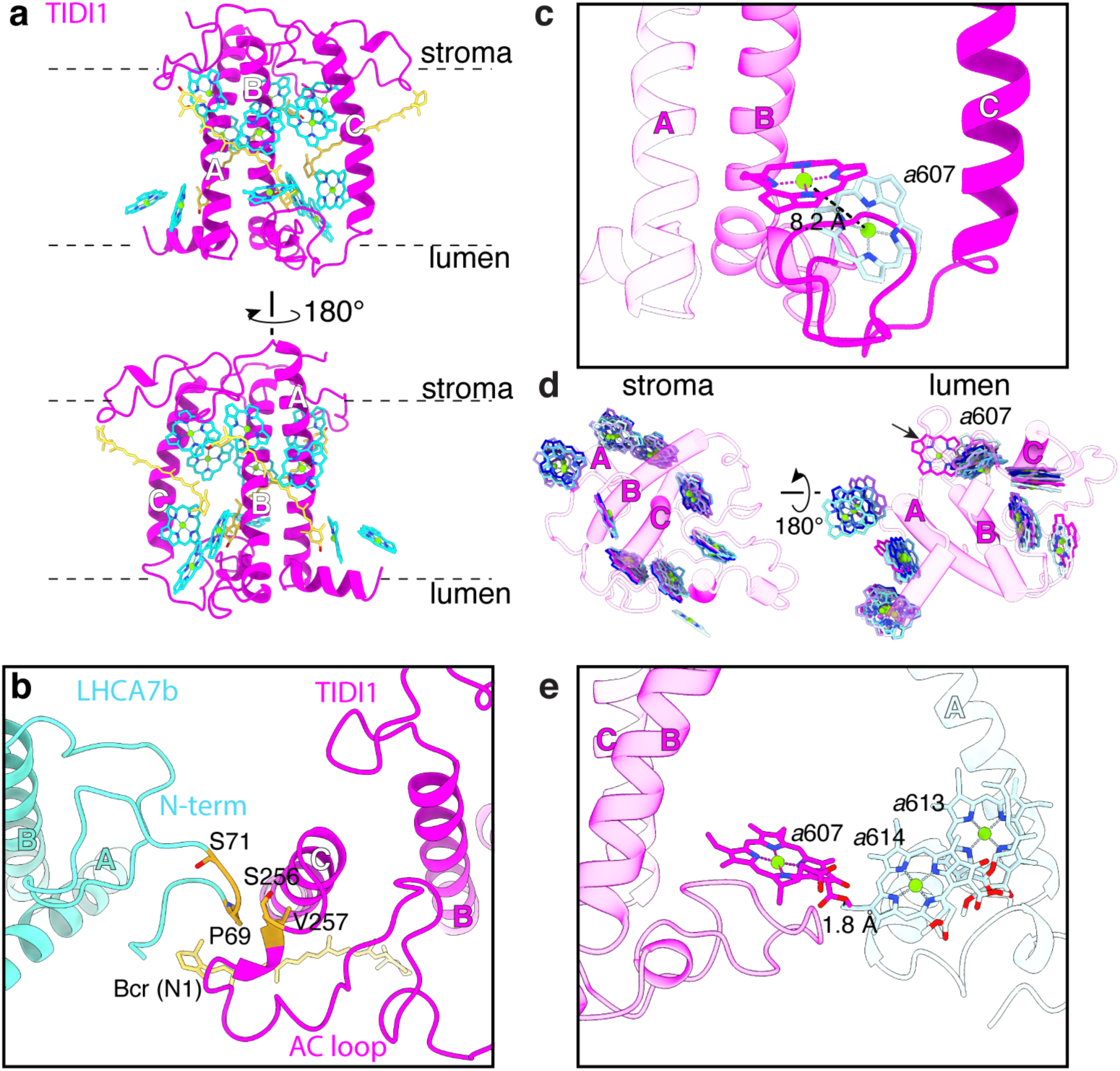
The extended BC loop of *Dt*TIDI1 differently positions Chl *a*607. **a**, Cartoon representation of *Dt*TIDI1. Chls are shown as cyan stick representations, with the central Mg atoms shown as spheres and numbered according to the conserved sites in spinach LHCII (PDB code 1RWT). Carotenoids are shown as yellow-colored stick representations. **b**, Interactions between *Dt*TIDI1 and *Dt*LHCA7b in the outer LHCI tetramer. Residues involved in the interaction are highlighted in orange. **c**, Comparison of *a*607_TIDI1_ against position of *a*607_LHCA3_ from *D. tertiolecta* from the lumenal side. Distance between molecules are shown as dashed lines. **d**, Comparison of *Dt*TIDI1 Chl positions against Chl positions of *Dt*LHCA1, *Dt*LHCA2, *Dt*LHCA3, *Dt*LHCA7, *Dt*LHCA8, and *Dt*LHCA9 from *D. tertiolecta* on the stromal (left) and lumenal side (right). **e**, Side view of *D. tertiolecta a*607_TIDI1_ (magenta) and its distance shown as to *a*614_LHCA3_ (light blue).

**Extended Data Fig. 21.**
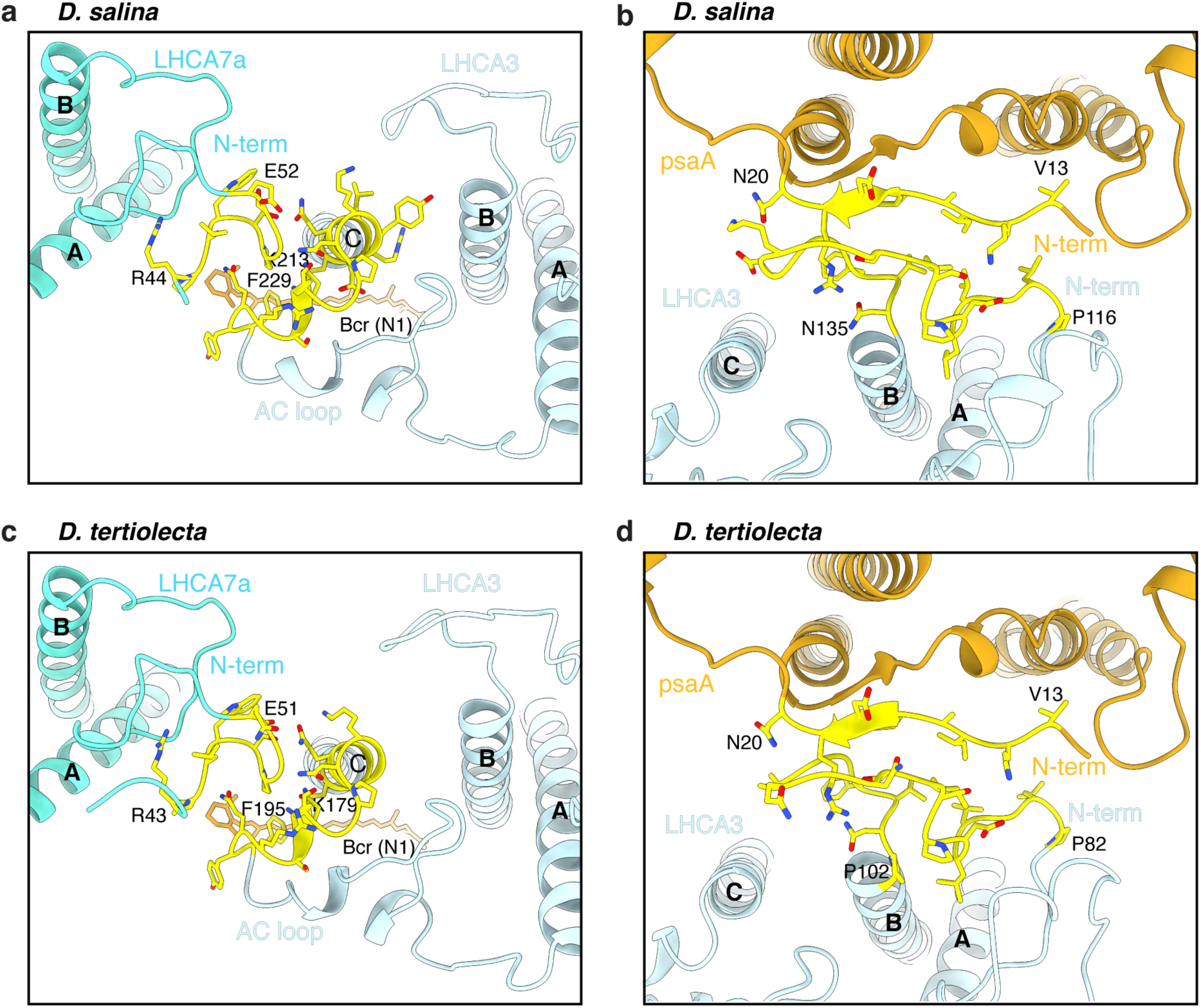
Interactions between LHCA3 and either LHCA7a or PsaA. **a-d**, Interactions between LHCA3 and LHCA7a in the inner LHCI tetramer (**a, c**) or between LHCA3 and PsaA in the PSI core (**b, d**) in either *Ds*PSI-LHCI_2_ supercomplex or *Dt*PSI-LHCI_2_ supercomplex. Residues involved in the interaction are shown as yellow stick representations.

**Extended Data Fig. 22.**
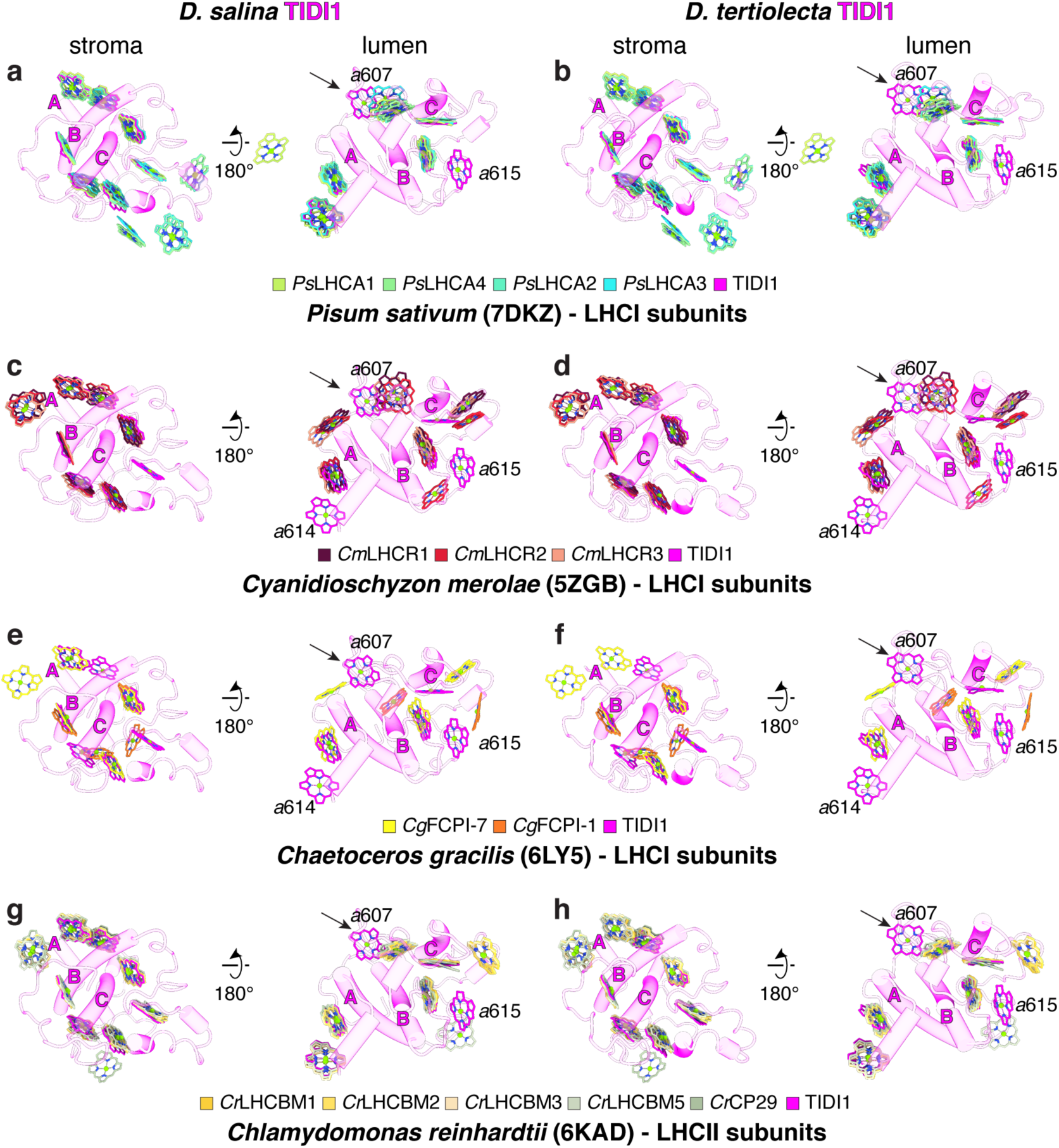
Chlorophyll 607 is unique to *Dunaliella* TIDI1 across the green lineage. Comparison of chlorophyll arrangement in *Ds*TIDI1 is shown in magenta and higher plants (*P. sativum*, pea) (**a**,**b**), red algae (*Cyan. merolae*) (**c**,**d**), diatoms (*Chae. gracilis*) (**e**,**f**), and LHCII subunits (*Chla. reinhardtii*) (**g**,**h**) at the stromal layer (left) and at the lumenal layer (right) for *D. salina* and *D. tertiolecta*, respectively. Nomenclature of chlorophylls is based on LHCII structure. Transmembrane helices are labeled in all of the panels.

**Extended Data Fig. 23.**
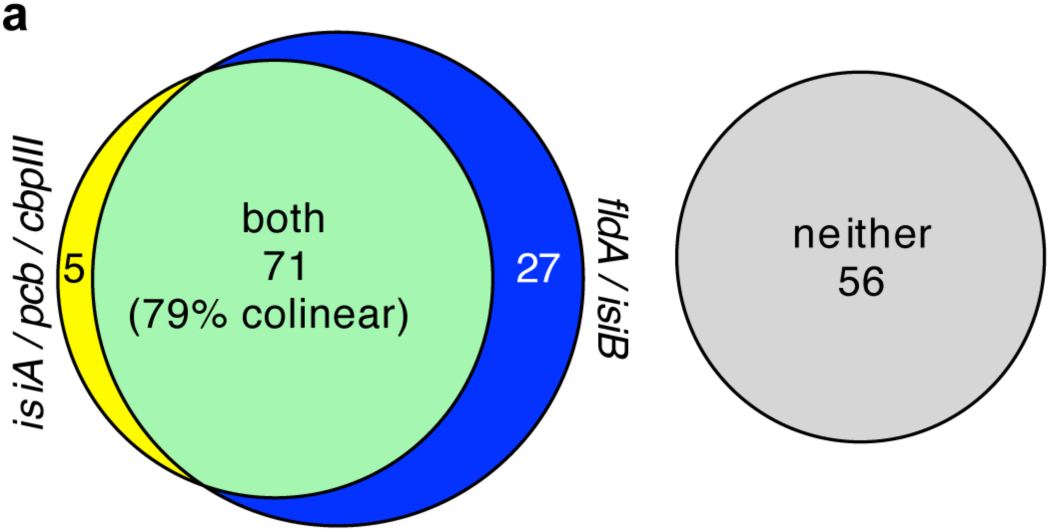
The co-occurrence of *isiA* and *fldA* genes in Cyanobacteria. **a**,159 high-quality genome assemblies in the Cyanobacteriota lineage were surveyed for the presence of *isiA* (also called *cbp* and indistinguishable from *pcb*) and *fldA (*also called *isiB)* genes. Here, Euler plots show the percentage of species with one or more copies of *isiA*, with one or more copies of *fldA*, with both, or with neither. *isiA* and *fldA* genes were considered colinear if they were within 5 kb of each other on the same strand of the same contig in a species’ genome. Supporting data are provided in Extended Dataset S3.

**Extended Data Fig. 24.**
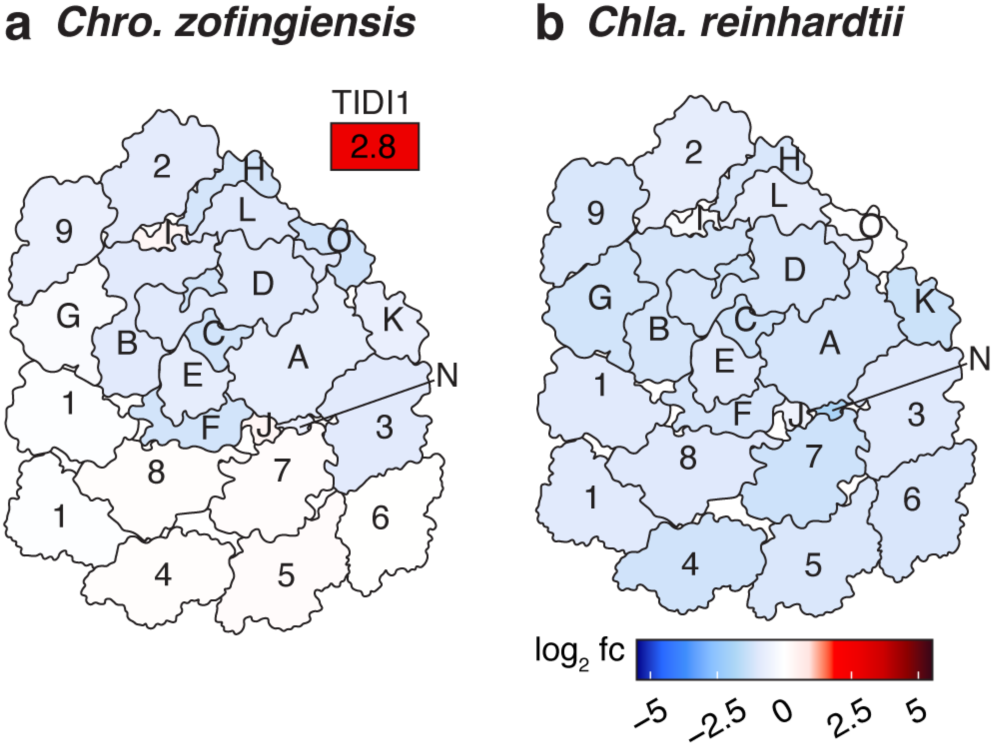
Fe-starvation responses of PSI and LHCI subunits in diverse green algae. **a**,**b**, The surface of PSI-LHCI supercomplexes colored by their log_2_ fold changes (fc) of the Fe-starved condition compared to the Fe-replete condition (red, increase; blue, decrease) for phototrophically grown *Chro. zofingiensis* (arrangement based on PDB:7D0J) (**a**) and *Chla. reinhardtii* (PDB:7D0J) (**b**).

**Extended Table 1.**
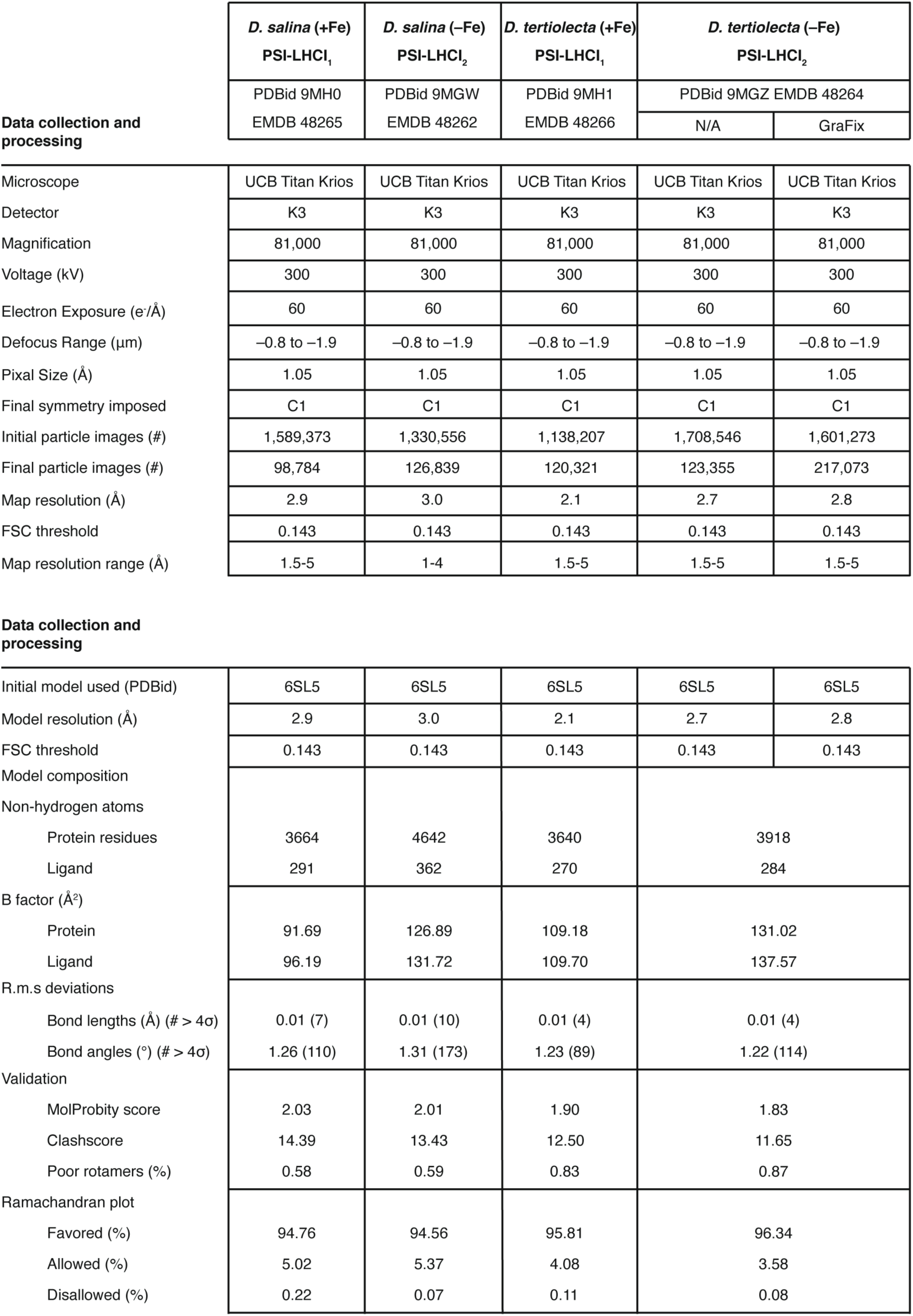
Data collection and processing.

**Extended Table 2.** Cofactors in each subunit of the Fe-replete *Dunaliella* spp. PSI-LHCl_1_ supercomplex.

| Species | Subunit | Traced residues | Chls | Cars | Lipids | Others |
| --- | --- | --- | --- | --- | --- | --- |
| *D. salina* | PsaA | 740 (12-751) | 42 *a*, 1 *a'* | 7 βCR | 3 LHG, 2 LMU, 1 OCD | 1 PQ, SF4 |
| | PsaB | 732 (3-734) | 41 *a* | 7 βCR | 1 DGD, 3 LHG, 1 SQD | 1 PQ |
|  | PsaC | 80 (2-81) |  |  |  | 2 SF4 |
|  | PsaD | 143 (60-202) |  |  |  |  |
|  | PsaE | 64 (60-123) |  |  |  |  |
| | PsaF | 165 (68-232) | 4 *a* | 2 βCR | 2 LHG, 3 PTY | |
| | PsaG | 105 (37-141) | 3 *a* | 1 βCR | 2 LHG | |
| | PsaH | 97 (69-133) | 2 *a* | 1 βCR | | |
| | PsaI | 41 (68-108) | | 2 βCR | | |
| | PsaJ | 41(1-41) | 1 *a* | 2 βCR | 1 LHG | |
| | PsaK | 83 (41-123) | 4 *a* | 1 βCR | | |
| | PsaL | 157 (47-203) | 4 *a* | 2 βCR | 1 4RF, 1 DGD, 1 PTY | |
| | LHCA1a | 197 (32-228) | 12 *a*, 2 *b* | 1 βCR, 1 XAT, 1 LUT | 2 LHG, 2 LMG | |
| | LHCA7a | 209 (63-271) | 10 *a*, 4 *b* | 1 βCR, 1 XAT, 1 LUT | 1 DGD, 3 LHG, 2 LMG, 1 PTY | |
| | LHCA8a | 226 (28-253) | 11 *a*, 3 *b* | 1 βCR, 1 XAT, 1 LUT | 2 LHG, 1 LMK, 2 PTY | |
| | LHCA3 | 208 (112-319) | 13 *a*, 1 *b* | 3 βCR, 1 XAT, 1 LUT | 1 DGD, 1 LHG, 2 LMG, 1 PTY, 2 SQD | |
|  | LHCA2 | 222 (41-262) | 11 *a*, 2 *b* | 4 LUT | 3 LHG, 2 PTY, 1 SQD, 1 3PH |  |
|  | LHCA9 | 187 (35-221) | 11 *a*, 1 *b* | 2 XAT, 1 LUT | 1 LHG, 1 LMU, 1 PTY, 1 SQD |  |
| *D. tertiolecta* | PsaA | 740 (12-751) | 42 *a*, 1 *a'* | 7 βCR | 3 LHG, 2 LMU, 1 PTY | 1 PQ, SF4 |
| | PsaB | 734 (2-735) | 41 *a* | 7 βCR | 1 DGD, 2 LHG, 1 LMU, 2 SQD | 1 PQ |
|  | PsaC | 80 (2-81) |  |  |  | 2 SF4 |
|  | PsaD | 143 (69-211) |  |  |  |  |
|  | PsaE | 68 (62-129) |  |  |  |  |
| | PsaF | 165 (78-242) | 4 *a* | 2 βCR | 1 LMG, 1 SQD, 1 PTY | |
| | PsaG | 105(36-140) | 3 *a* | 1 βCR | 1 LMG | |
|  | PsaH | 47 (70-116) | 1 *a* |  |  |  |
| | PsaI | 35 (15-49) | | 2 βCR | | |
| | PsaJ | 41(1-41) | 1 *a* | 2 βCR | 1 LMU | |
| | PsaK | 83 (41-123) | 4 *a* | 1 βCR | | |
| | PsaL | 138 (56-193) | 2 *a* | 1 βCR | | |
| | LHCA1a | 197 (32-228) | 12 *a*, 2 *b* | 1 βCR, 1 XAT, 1 LUT | 1 LHG | |
| | LHCA7a | 217 (39-255) | 10 *a*, 4 *b* | 1 βCR, 1 XAT, 1 LUT | 3 LHG, 1 DGD, 1 SQD | |
| | LHCA8a | 226 (28-253) | 11 *a*, 3 *b* | 1 βCR, 1 XAT, 1 LUT | 3 LHG, 1 LMG, 1 LMU, 1 LMK, 1 PTY | |
| | LHCA3 | 223 (62-284) | 13 *a*, 1 *b* | 3 βCR, 1 XAT, 1 LUT | 2 LMG, 2 SQD, 2 PTY | |
|  | LHCA2 | 222 (39-260) | 11 *a*, 2 *b* | 2 LUT | 2 LHG |  |
|  | LHCA9 | 187 (35-221) | 11 *a*, 1 *b* | 2 XAT, 1 LUT | 2 LHG, 1 LMU, 1 PTY |  |

DGD, digalactosyldiacyl glycerol; LHG, dipalmitoyl phosphatidylglycerol; LMU, dodecyl- α-D-maltoside; PTY, phosphatidyl glycerol; SQD, sulfoquinovosyldiacyl glycerol; 3PH, phosphatidic acid; 4RF, tripalmitoylglycerol; OCD, octadecanal; PQN, phylloquinone; Chl, chlorophyll; Cars, carotenoids; SF4, sulfur-iron cluster

**Extended Table 3.** Cofactors in each subunit of the Fe-starved *D. salina* PSI-LHCl_2_ supercomplex.

| Species | Subunit | Traced residues | Chls | Cars | Lipids | Others |
| --- | --- | --- | --- | --- | --- | --- |
| *D. salina* | PsaA | 737 (12-748) | 43 *a*, 1 *a'* | 7 βCR | 3 LHG, 2 LMU, | 1 PQ, SF4 |
|  | PsaB | 732 (3-734) | 41 *a* | 6 βCR | 1 DGD, 2 LHG, 2 PTY | 1 PQ |
|  | PsaC | 80 (2-81) |  |  |  | 2 SF4 |
|  | PsaD | 143 (60-202) |  |  |  |  |
|  | PsaE | 64 (60-123) |  |  |  |  |
|  | PsaF | 165 (68-232) | 5 *a* | 2 βCR | 1 LMG, 1 PTY |  |
|  | **PsaG** | 105 (37-141) | 3 *a* | 1 βCR | 1 LMG |  |
|  | **PsaH** | 98 (36-133) | 2 *a* | 1 βCR |  |  |
|  | PsaI | 41 (68-108) |  | 2 βCR |  |  |
|  | PsaJ | 41(1-41) | 1 *a* | 2 βCR |  |  |
|  | PsaK | 83 (41-123) | 4 *a* | 1 βCR |  |  |
|  | PsaL | 160 (44-203) | 4 *a* | 2 βCR | 1 PTY, 1 4RF |  |
|  | **PsaO** | 90 (40-129) | 3 *a* | 1 βCR, 1 LUT |  |  |
|  | LHCA1a | 197 (32-228) | 12 *a*, 2 *b* | 1 βCR, 1 XAT, 1 LUT | 2 LHG, 1 LMG, 1 LMU, 1 3PH |  |
|  | LHCA7a | 209 (43-251) | 10 *a*, 4 *b* | 1 βCR, 1 XAT, 1 LUT | 1 DGD, 4 LHG, 1 LMG, 1 PTY |  |
|  | LHCA8a | 226 (28-253) | 11 *a*, 3 *b* | 1 βCR, 1 XAT, 1 LUT | 1 LHG, 1 LMK, 2 PTY |  |
|  | LHCA3 | 230 (90-319) | 13 *a*, 1 *b* | 3 βCR, 1 XAT, 1 LUT | 2 DGD, 1 LHG, 2 LMG, 1 PTY, 1 SQD |  |
|  | LHCA1b | 196 (32-227) | 12 *a*, 2 *b* | 1 βCR, 1 XAT, 1 LUT | 2 LHG, 1 PTY |  |
|  | LHCA7b | 209 (43-251) | 10 *a*, 4 *b* | 1 βCR, 1 XAT, 1 LUT | 1 LHG, 3 PTY |  |
|  | LHCA8b | 223 (29-251) | 11 *a*, 3 *b* | 1 βCR, 1 XAT, 1 LUT |  |  |
|  | TIDI1 | 205 (148-352) | 12 *a*, 1 *b* | 2 βCR, 1 XAT, 1 LUT | 1 LHG |  |
|  | **LHCA2** | 221 (41-261) | 11 *a*, 2 *b* | 4 LUT | 2 LHG, 2 PTY, 1 SQD, 1 3PH |  |
|  | **LHCA9** | 187 (35-221) | 11 *a*, 1 *b* | 2 XAT, 1 LUT | 2 LHG, 1 LMU, 1 PTY |  |

DGD, digalactosyldiacyl glycerol; LHG, dipalmitoyl phosphatidylglycerol; LMU, dodecyl-α-d-maltoside; PTY, phosphatidyl glycerol; SQD, sulfoquinovosyldiacyl glycerol; 3PH, phosphatidic acid; 4RF, tripalmitoylglycerol; PQN, phylloquinone; Chl, chlorophyll; Cars, carotenoids; SF4, sulfur-iron cluster; Red color denotes subunits absent in the *D. tertiolecta* PSI-LHCI<sub>2</sub> supercomplex

**Extended Table 4.** Cofactors in each subunit of the Fe-starved *D. tertiolecta* PSI-LHCl_2_ supercomplex.

| Species | Subunit | Traced residues | Chls | Cars | Lipids | Others |
| --- | --- | --- | --- | --- | --- | --- |
| *D. tertiolecta* | PsaA | 740 (12-751) | 42 *a*, 1 *a'* | 7 βCR | 2 LHG, 2 LMU | 1 PQ, SF4 |
|  | PsaB | 734 (2-735) | 42 *a* | 6 βCR | 1 DGD | 1 PQ |
|  | PsaC | 80 (2-81) |  |  |  | 2 SF4 |
|  | PsaD | 143 (51-193) |  |  |  |  |
|  | PsaE | 67 (43-109) |  |  |  |  |
|  | PsaF | 165 (63-227) | 4 *a* | 2 βCR | 1 LMG |  |
|  | PsaI | 35 (15-49) |  | 2 βCR |  |  |
|  | PsaJ | 41(1-41) | 1 *a* | 2 βCR |  |  |
|  | PsaK | 83 (41-123) | 4 *a* | 1 βCR |  |  |
|  | PsaL | 138 (56-193) | 2 *a* | 1 βCR |  |  |
|  | LHCA1a | 197 (32-228) | 12 *a*, 2 *b* | 1 βCR, 1 XAT, 1 LUT | 3 LHG, 2 PTY |  |
|  | LHCA7a | 217 (39-255) | 10 *a*, 4 *b* | 1 βCR, 1 XAT, 1 LUT | 1 DGD, 2 LHG |  |
|  | LHCA8a | 226 (28-253) | 11 *a*, 3 *b* | 1 βCR, 1 XAT, 1 LUT | 1 LHG, 1 LMK, 2 PTY |  |
|  | LHCA3 | 226 (58-283) | 13 *a*, 1 *b* | 3 βCR, 1 XAT, 1 LUT | 1 DGD, 2 LMG, 1 SQD |  |
|  | LHCA1b | 197 (32-228) | 12 *a*, 2 *b* | 1 βCR, 1 XAT, 1 LUT | 2 LHG |  |
|  | LHCA7b | 209 (42-250) | 10 *a*, 4 *b* | 1 βCR, 1 XAT, 1 LUT | 1 LHG |  |
|  | LHCA8b | 224 (28-251) | 11 *a*, 3 *b* | 1 βCR, 1 XAT, 1 LUT | 1 LHG |  |
|  | TIDI1 | 210 (130-339) | 11 *a*, 1 *b* | 1 βCR, 1 XAT, 1 LUT |  |  |

DGD, digalactosyldiacyl glycerol; LHG, dipalmitoyl phosphatidylglycerol; LMU, dodecyl-alpha-d-maltoside;PTY, phosphatidyl glycerol; SQD, sulfoquinovosyldiacyl glycerol; PQN, phylloquinone; Chl, chlorophyll; Cars, carotenoids; SF4, sulfur-iron cluster

## Notes

### Competing Interest Statement

The authors have declared no competing interest.

